# An objective criterion to evaluate sequence-similarity networks helps in dividing the protein family sequence space

**DOI:** 10.1101/2022.04.19.488343

**Authors:** B. V. H. Hornung, N. Terrapon

**Affiliations:** Aix Marseille Universite, CNRS, UMR 7257 AFMB, Marseille, France; INRAE, USC 1408 AFMB, Marseille, France

**Keywords:** CAZy, carbohydrate active enzymes, sequence-based protein classification, protein subfamilies, SSN, sequence similarity network

## Abstract

The deluge of genomic data raises various challenges for computational protein annotation. The definition of superfamilies, based on conserved folds, or of families, showing more recent homology signatures, allow a first categorization of the sequence space. However, for precise functional annotation or the identification of the unexplored parts within a family, a division into subfamilies is essential. As curators of an expert database, the Carbohydrate Active Enzymes database (CAZy), we began, more than 15 years ago, to manually define subfamilies based on phylogeny reconstruction. However, facing the increasing amount of sequence and functional data, we required more scalable and reproducible methods. The recently popularized sequence similarity networks (SSNs), allows to cope with very large families and computation of many subfamily schemes. Still, the choice of the optimal SSN subfamily scheme only relies on expert knowledge so far, without any data-driven guidance from within the network. In this study, we therefore decided to investigate several network properties to determine a criterion which can be used by curators to evaluate the quality of subfamily assignments. The performance of the closeness centrality criterion, a network property to indicate the connectedness within the network, shows high similarity to the decisions of expert curators from eight distinct protein families. Closeness centrality also suggests that in some cases multiple levels of subfamilies could be possible, depending on the granularity of the research question, while it indicates when no subfamily emerged in some family evolution. We finally used closeness centrality to create subfamilies in four families of the CAZy database, providing a finer functional annotation and highlighting subfamilies without biochemically characterized members for potential future discoveries.

**Author Summary:** Proteins perform a lot of functions within living cells. To determining their broad function, we group similar amino-acid sequences into families as their shared ancestry argue for shared functionality. That’s what we do in the CAZy database, which covers >300 Cazbohydrate-Active enZyme families nowadays. However, we need to divide families into subfamilies to provide finer readibility into (meta)genomes and guide biochemists towards unexplored regions of the sequence space. We recently used the popularized Sequence Similarity Networks (SSN) to delineate subfamilies in the large GH16 family, but had to entirely rely on expert knowledge to evaluate and take the final decision until now, which is not scalable, not eough automated and less reproducible. To accelerate the construction of protein subfamilies from sequence similarity networks, we present here an investigation of different network properties, to use as indicators for optimal subfamily divisions. The closeness centrality criterion performed best on artificial data, and recapitulates the decisions of expert curators. We used this criterion to divide four more CAZy families into subfamilies, showed that for others no subfamilies exist.

We are therefore able to create new protein subfamilies faster and with more reliability.

## Introduction

The amount of newly produced genomic data increases exponentially. This leads to various challenges in how to treat this data, including how to store and analyze it. Another of these challenges is the functional annotation of such data. Ideally, functional annotation is obtained after the experimental demonstration of protein activity, but due to the deluge of data this is impossible (1,2). Computational annotation is performed instead in the vast majority of cases. Many methods are available to annotate genomes (2), and most rely on the assignment of sequences to groups of homologs. This is followed by the transfer of the annotation of experimentally characterized proteins to non-characterized group members. In this context, several levels of granularity can be considered from superfamilies to subfamilies. Superfamilies group many remote homologs that display a similar fold but can largely vary in their molecular functions/specificities, only allowing to have a very general annotation but for many proteins (3). In contrast, subfamilies tend to group very few but closely-related sequences, offering a more reliable/precise annotation but only when at least one member has been biochemically characterized. For the delineation of subfamilies, sequence similarity networks (SSNs) have become increasingly popular (4–6). SSNs consist in running pairwise comparisons (typically BLASTp) between all family members, and select all matching pairs satisfying a given criterion (typically an e-value cutoff, sometimes with coverage requirements) to draw a network in which the connected components would be subfamilies. The main advantage of SSNs compared to phylogeny is their ability to cope with large families for which the alignment computation and results are problematic. Their main limitation is that (i) they require the exploration of the many possible cutoffs and (ii) the critical step for the determination of the optimal solution fully relies on human expertise. While expert knowledge is arguably one of the very principles of science, it lacks the scalability required to deal with nowaday’s amount of data and the reproducibility for sharing scientific advances. For experts, SSN tools do not necessarily produce human-friendly visualization of the results, making the selection of a suitable network, out of all possible SSNs, a real challenge. As curators of a specialist resource, the Carbohydrate-Active enZyme (CAZy) database, one of our objectives is to enhance the predictive power of our protein annotation, since our community relies on our annotation and expert knowledge to build hypothesis leading to future discoveries (7). Due to the large amount of CAZy families (currently >300) and the existence of very large and multifunctional families, there is an urgent need to make the subfamily investigations more performant, as well as reproducible. We therefore decided to investigate currently available methods and indicators from which the optimal SSN can easily and reproducibly be derived by expert curators. We investigated various clustering methods, dimensionality reduction techniques and various network properties, which indicate grouping and spread of nodes within the network. We applied these methods to artificial datasets, as well as to the recently published SSN of glycoside hydrolase family GH16 (8) and to various families from the structure function linkage database (9). We show that the *closeness centrality* criterion is a fast and relevant indicator for optimal subfamily delineations. It allows the curator to identify a few possibly optimal networks out of often more than a hundred produced in SSN computations. We furthermore show that in some cases multiple levels of subfamilies could be possible, whereas, in others, no subfamilies exist. After determining this as the appropriate method, we generated and analyzed the networks of nine CAZy families, accelerating our efforts in subfamily creation.

## Results

Our goal was to identify an objective criterion to determine optimal SSN e-value cutoffs, given that our assumption is that an optimal cutoff should maximize the number of connections within the subnetworks/subfamilies, without having any connections towards the other subnetworks/subfamilies. We initially investigated various network properties on artificial data and identified the *closeness centrality* as the best suited. This criterion was used to reanalyze already published data based on real families of protein sequences, showing overall good agreement. We further compared SSN results to clustering and dimensionality reduction (see supplementary results), and confirmed the better suitability of the SSNs. We therefore applied this criterion to guide the analysis of subfamilies for several CAZy families, glycoside hydrolase family GH19, GH51, GH45, GH54, GH55, GH68 and GH130 and the polysaccharide lyase family PL26.

### Closeness centrality is the best indicator on artificial data

Artificial data was generated (see Methods) to simulate a protein sequence family composed of at least one, or two, subfamilies. In real data, such family would appear as a large network at high e-value thresholds, as all members display a detectable similarity to all others, either directly or by transitivity. With the decrease of the e-value cutoff, edges successively disappear leading to the disconnection of single-nodes or subnetworks which should correspond to subfamilies (Figure 1).

**Figure 1:**
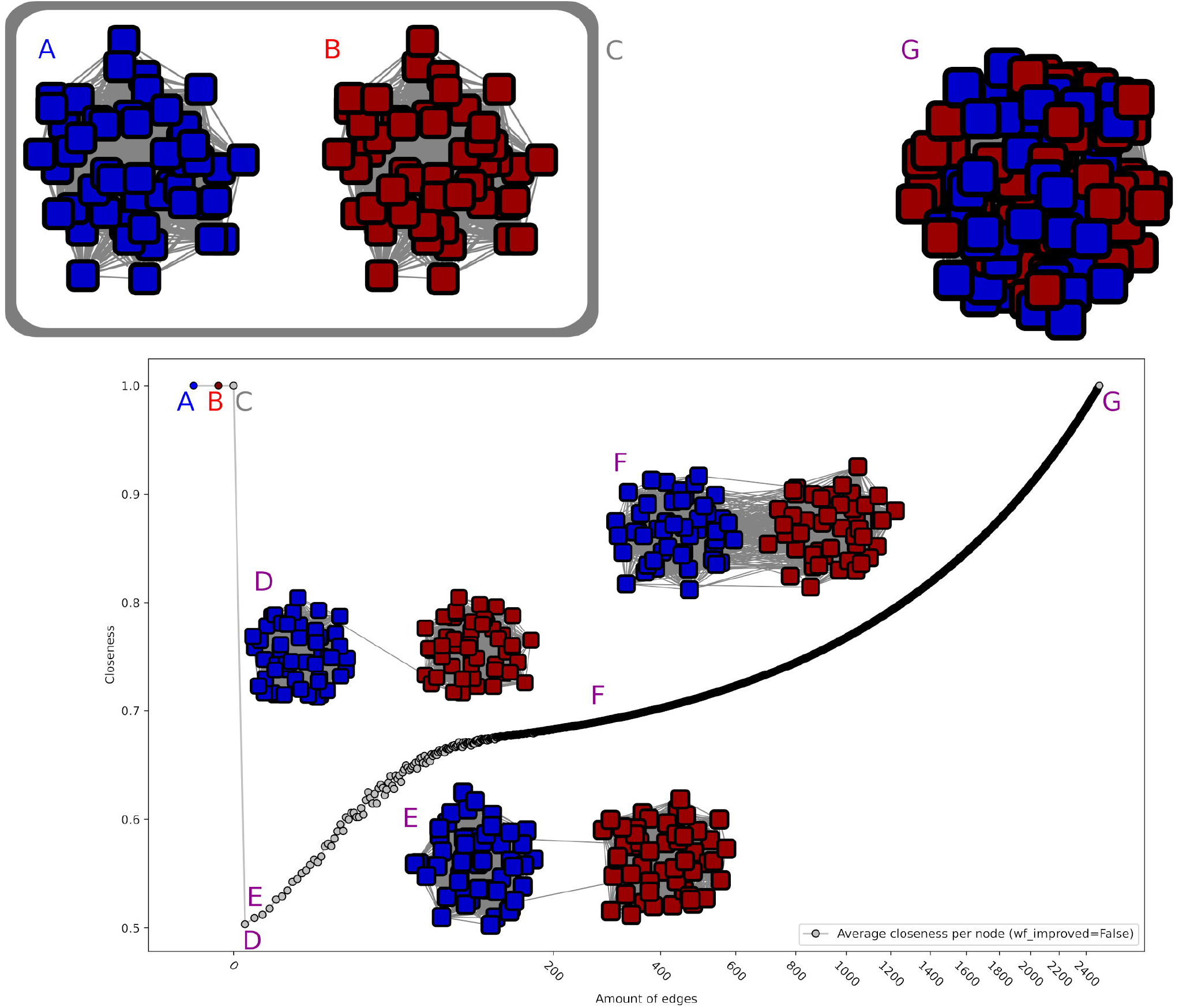
Illustration of closeness centrality behaviour on artificial data.Closeness values were averaged over all nodes using two perfect subnetworks of size 50, which sequentially get connected by edges. As it can be seen in A and B, each perfect subnetwork separately reaches a closeness centrality of 1, as well as the combined network C. In all three cases no new edges have been added yet. This value increases with each added edge, slighthy with one or two edges (D and E), to intermediate values (e.g. F, 200 edges), up again to a maximum value if all edges are connected at G. The critical point is between C and D. While only one component is present in D, it gets evaluated a lot lower than C, despite having more edges.

Our artificial data thus consisted in *network collections*. Each *collection* is designed to simulate the SSNs of a family for increasing e-value thresholds, one *network* corresponding to one threshold and containing more edges than the previous and less than the next. Hence all networks in the *collection* are constructed upon two identical subnetworks but could be distinguished by the number of edges between the subnetworks (interconnections but no intraconnections), from one to all possible connections. The different *collections* should simulate families with distinct evolutionary properties, from one (the main network without any subfamilies) to many subfamilies, and correspond to the combination of different types of subnetworks (from fully-connected to single-node subnetworks).

We investigated the behavior of several classical network properties like degree centrality, betweenness centrality and closeness centrality on the *artificial* data. The ideal property should reach its global optimum for a network, which maximizes the connections within its subnetworks, and shows no connections to other subnetworks. In our artificial setting, it should indicate as local optima both the networks consisting of (i) all connections between the two subnetworks and (ii) the two disconnected subnetworks. The desired behavior was obtained from *closeness centrality* (with *wf_improved* parameter set to False) (Figure 1). Especially this is interesting to note the partially logarithmic behavior of closeness centrality for the lowest number of connections, which thus scores extremely low. *Betweenness centrality* (and its variations, listed in the methods) showed a similar behavior in some network combinations, but was discarded after investigating its behavior in combination with single-node networks.

Indeed, *Betweeness centrality* weights singles as perfect networks, which is not a desired behavior, since this would favor strongly disconnected networks. A network consisting out of only singles would be evaluated to be as good as a perfect network, which is not what we were looking for. All other criteria behaved either similarly or identical to one of these (one similar to closeness centrality, four similar to betweenness, where betweenness centrality source, load centrality and betweenness behave identical), were monotonous in their behavior (e.g. two perfect networks with a single connection was evaluated better than a single perfect network; five in total), or not quantitative enough (e.g. asynchronous label propagation indicating one or two subnetworks; three criteria). The remaining four other criteria were (i) the eigenvector centrality and the clustering coefficient, where the former showed parabola-like behavior and the latter an inverted parabola behavior, therefore scoring better with less and more edges than with a medium number of edges, which is the opposite of the desired behavior; (ii) edge load centrality and global reaching centrality, which both partially showed a bimodal distribution. This means that with these both algorithms either both less and more edges would be favored or disfavored. An overview over all behaviors on two networks with a size of 50 can be seen in supplementary Figure S3.

Based on artificial data results, we therefore opted to use *closeness centrality (*with wf_improved set to False), applied to each subnetwork, as criterion for further evaluation.

### *Closeness centrality* supports findings by expert curators

To confirm the relevance of closeness centrality, it was applied to determine the optimal cutoffs in published SSN analysis by several independent research groups, based on real protein family datasets, to compare with the cutoff determined by expert curators. We created the SSNs of one Interpro family, three SFLD families, and two CAZy glycoside hydrolase families.

We first compared the closeness centrality performance to an analysis that used the popular SSN tool EFI-EST (5). We selected one among its most recent citations in PubMed (September 2020), and with easily reproducible methods (10). In this publication about the domain IPR037434, Travis *et al*. identified three relevant cutoffs including 10^−55^ corresponding to 5 subfamilies with more than 3 members. We generated SSNs from 10^−5^ to 10^−100^ (steps of 10^−5^), on which *closeness centrality* computations show a local optimum for an e-value of 10^−60^ (difference possibly influenced by the increase in the database size; 228 sequences in Travis et al., 376 in this investigation), which corresponds to 10 families with more than 3 members (51 subfamilies total). A consensus probably resides between those two.

For SFLD family 159 of tautomerases (11), the authors concluded first on an e-value of 10^−11^ for subfamily assignment, but due to the great diversity in their sequences also further investigated an e-value of 10^−18^ to define 18 subfamilies. The closeness centrality computations show local optima at e-values of 10^−9^ and 10^−18^, the latter displaying 15 subfamilies with >100 members.

However, an absolute maximum was obtained at 10^−34^, although this would result in 12 more subfamilies (with >100 members), and more than 2000 smaller networks. For SFLD subfamily 19 of glutathione transferases (12), two cutoffs were identified by the authors, 10^−13^ and 10^−25^, the second level allowing more detailed inspections. The closeness centrality computations for this family peak at 10^−14^ and 10^−29^, with an absolute maximum at 10^−44^. For SFLD family 122 of nitroreductases, the published cutoff of 10^−18^ (13) also corroborates with the first local optimum of closeness centrality computations at 10^−19^. The global optimum was still not reached at 10^−69^, where we stopped our computations.

For our own recent SSN analysis of family GH16 (8), an e-value of 10^−55^ was published (SSNs generated in increments of 10^−5^), while closeness centrality revealed a peak at 10^−50^. Similarly, for GH130 (14), the authors determined the separation at 10^−70^, corresponding to 15 relevant subfamilies (>20 members). Closeness centrality reaches a peak at 10^−75^, resulting in 16 such relevant subfamilies. The subfamilies in CAZy are created according to the numbering of (14), with the omission of families purely derived from metagenomic data (UC4 and UC6) replaced by additional subfamily we discovered (GH130_10 and GH130_12).

### *Closeness centrality* accurately points to suitable novel subfamily schemes

Closeness centrality was therefore applied to analyze SSNs generated for several CAZy families for which no subfamily scheme was designed yet. The family selection was based on potential interest for on-going projects, as well as on the diversity and total number of functional annotations available. Functional annotations are essential indicators to decide which scheme will be the most relevant and durable, among the several optimal options suggested by closeness centrality.

We first investigated the small GH55 family (1052 sequences after fragment filtering). This family also display low functional diversity as all characterized members breakdown b-1,3-glucans (mainly tested on laminarin), with the activity only differing by the mode of action: exo (EC 3.2.1.58), endo (EC 3.2.1.39), or undetermined (appearing in CAZy as laminarin-degrading enzymes, EC 3.2.1.-). Many characterized enzymes were described by Bianchetti and collaborators (15) who hypothesized that most members should be exo-acting. In their analysis of 177 GH55 proteins, the reconstructed phylogeny suggests three clades: a fungal, a bacterial and a divergent (bacterial) clade. The closeness centrality values for the generated SSNs indicated a local optimum at 10^−12^ and the global optimum, close to maximum, at 10^−29^ (Figure 2). At 10^−12^, two subfamilies emerged, one corresponding to the bacterial clade, the second gathering the fungal and “divergent” clades previously described. At 10^−29^, the fungal and divergent subfamilies split, resulting in a nearly perfect closeness centrality value. The two distinct modes of action did not segregate among the three subfamilies, given both are retrieved in the bacterial and in the fungal subfamilies, while the divergent subfamily does not include any characterized representative. As a consequence, GH55 will be divided into subfamilies 1 to 3, by ascending size order, which correspond respectively to the bacterial, fungal and divergent subfamilies.

**Figure 2:**
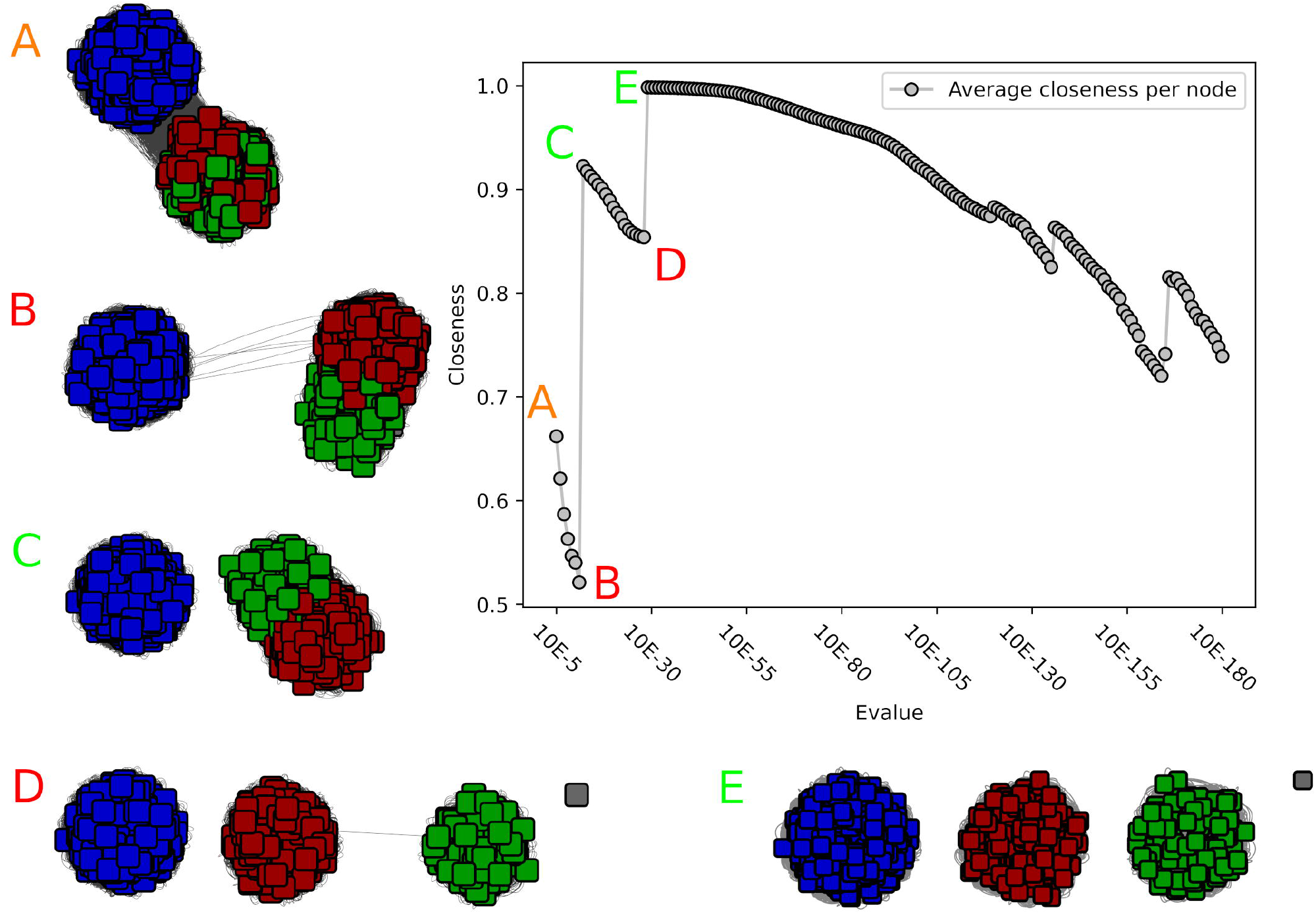
Closeness value and corresponding networks of the GH55 SSN analysis. A) At 10^−5^ the network is still a single network, and subnetworks become visible. The closeness value is in a medium range. B) The closeness value reaches its lowest point at 10^−11^. The two subnetworks within GH55 are only sparsely connected. C) At 10^−12^ the two subnetworks are disconnected, and the closeness value increases. D) The closeness value reaches another low at 10^−28^, and two other subnetworks are clearly visible, only sparsely connected with a single edge. A single disconnected node has emerged. E) At 10^−29^ three nearly perfect subnetworks are visible, and the closeness value is nearly perfect with a value of 0.99.

### Multiple levels of subfamilies are possible

We investigated another small CAZy family, GH68 (1510 sequences after filtering). Three enzymatic activities were reported in this transglycosidase family (i.e. able to not only breakdown but also assemble glycans depending on the concentration of the substrate and product of the reaction): invertase (b-fructofuranosidase; EC 3.2.1.26), inulosucrase (sucrose:fructan 1-fructosyltransferase; EC 2.4.1.9) and levansucrase (sucrose:fructan 6-fructosyltransferase; EC 2.4.1.10). In this family, closeness computations never reached the initial maximum of the fully-connected network (10^−5^). However, several local optima could be considered notably at 10^−51^, 10^−124^ and 10^−154^ corresponding to closeness centrality values of 0.93, 0.95 and 0.96 (Figure 3 C, G and H). At 10^−51^, the family would be divided into two subfamilies of similar size. At 10^−124^, both former subfamilies get split into two (all having more than 150 members) while only one subfamily with >20 members emerge (plus 17 tiny groups including 5 singletons). At 10^−154^, two more large groups emerge from the former four families.

**Figure 3:**
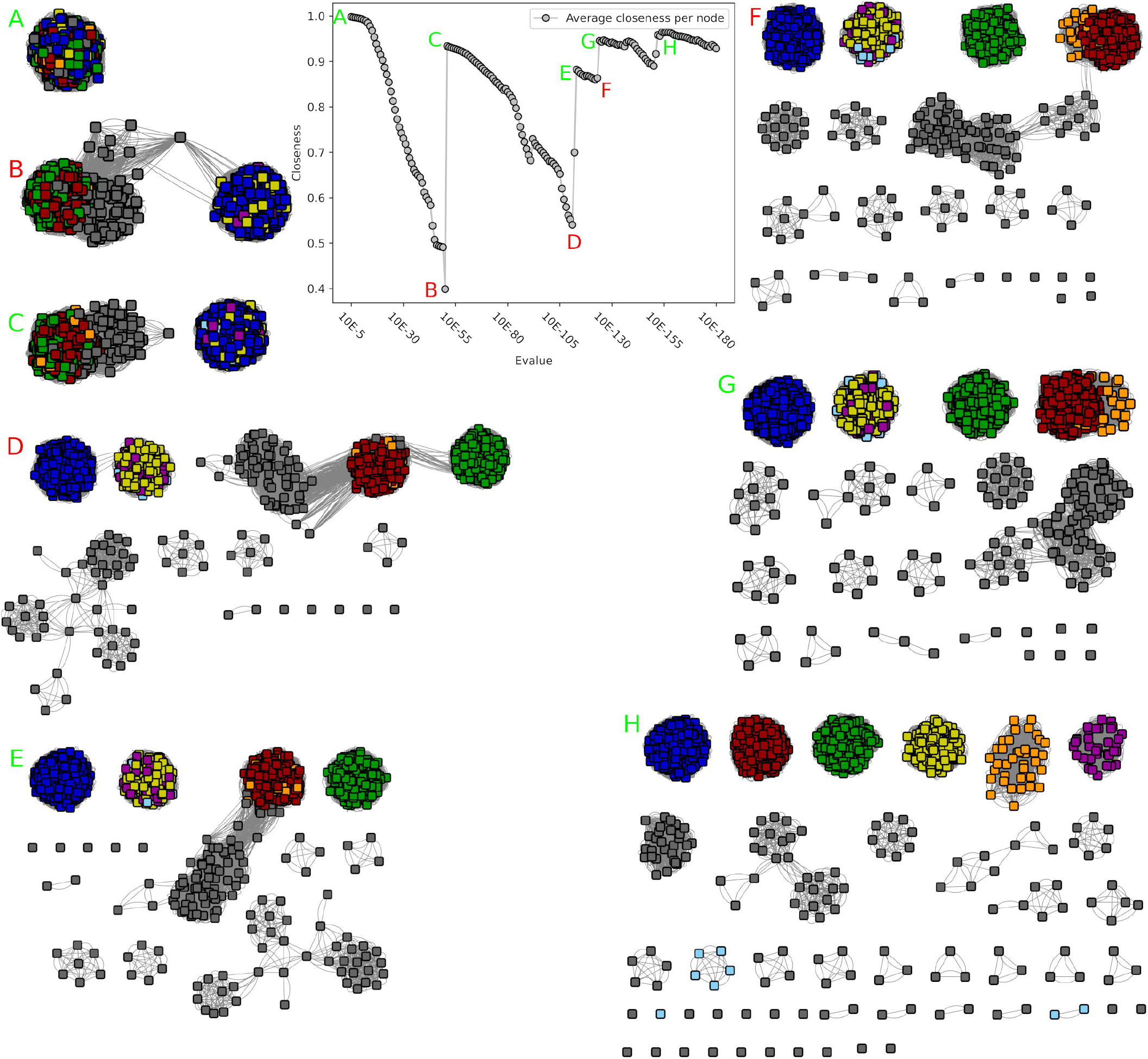
Closeness value and corresponding networks of the GH68 SSN analysis. Unlike in the GH55 analysis, the best network is less well determinable. A) At 10^−5^ the network is one solid component, and no subnetworks are visible. B) at 10^−50^ the closeness value is lowest, and two sparsely connected networks are visible. C) At 10^−51^ these two networks are disconnected, and the closeness value increases. D) to H) show the closeness values and networks at 10^−111^, 10^−113^, 10^−123^, 10^−124^ and 10^−154^ respectively. D and F show undesirable connections with sparely connected subnetworks, whereas E, G and H are alternative desirable network configurations which could be used to determine subfamilies.

At the functional level, at 10^−51^, the first subfamily gathers EC activities 2.4.1.9 and 2.4.1.10, whereas the second displays 2.4.1.9, 2.4.1.10 and 3.2.1.26 (including one protein with dual activities 2.4.1.9 and 2.4.1.10, as well as one with dual 2.4.1.10 and 3.2.1.26). At 10^−124^, each of the five main subfamilies include a characterized member, but most activities are shared these subfamilies, for example with 2.4.1.10 found in subfamilies 1 to 4; 2.4.1.9 in subfamilies 2, 4 and 5; and 3.2.1.26 in subfamilies 2, 3 and 5. At 10^−154^, 9 of the 19 subfamilies include a characterized member, five of which display a single activity. However, this activity is not unique but shared (see supplementary table 1), given the split is mostly driven by the taxonomy, as at 10^−124^. Therefore, GH68 will be divided into two subfamilies following 10^−51^ threshold, based on the absence of functional information gain, and the increasing number of unclassified sequences at lower optimal thresholds (none at 10^−51^ versus 74 and 96 at 10^−124^ and 10^−154^ resp., that is >6%).

We then analyzed the larger (>10,000 sequences after filtering) GH51 family. This family displays so far four different EC numbers on a total of 83 characterized enzymes, 76 sharing the same a-L-arabinofuranosidase activity (EC 3.2.1.55), five with an endo-b-1,4-glucanase activity (EC 3.2.1.4) and two bifunctional enzymes: an a-L-arabinofuranosidase/xylan 1,4-b-xylosidase (EC 3.2.1.55 and EC 3.2.1.37) and an endo-b1,4-glucanase/endo-b-1,4-xylanase (EC 3.2.1.4 and EC 3.2.1.8). The *closeness centrality* criterion showed again multiple possible optimal SSNs, at 10^−58^, 10^−91^ and 10^−147^, corresponding to closeness values of 0.71, 0.67 and 0.79 that are lower than previously analyzed families (Figure 4 A, C and D)). At 10^−58^, the network divides into two large subnetworks (5712 and 3960 sequences) and one small network (114 sequences), hereafter referred as to subfamilies 1 to 3, and 55 smaller subnetworks (<100 sequences). Both subfamilies 1 and 2 contain enzymes with EC 3.2.1.55 activity, and subfamily 2 also contained the bifunctional enzyme EC 3.2.1.55 and 3.2.1.37. Proteins in subfamily 2 often display a small conserved domain of unknown function (annotated in the Gene3D superfamily 2.60.120.260) in the N-terminal region, unlike the enzymes in subfamily 1, further supporting the split between these groups. Subfamily 3 contains all five characterized enzymes with the EC number 3.2.1.4, including the bifunctional enzyme with EC 3.2.1.4/3.2.1.8. Most enzymes in subfamily 3 also contain a CBM domain from families known for binding cellulose (CBM2, CBM11, CBM30), which are not present in subfamilies 1 and 2. A small subfamily (55 sequences) also showed a different domain architecture as most members contain a CBM66 module, and some are fused to a GH127 enzyme. These features are not seen in any of the other subfamilies.

**Figure 4:**
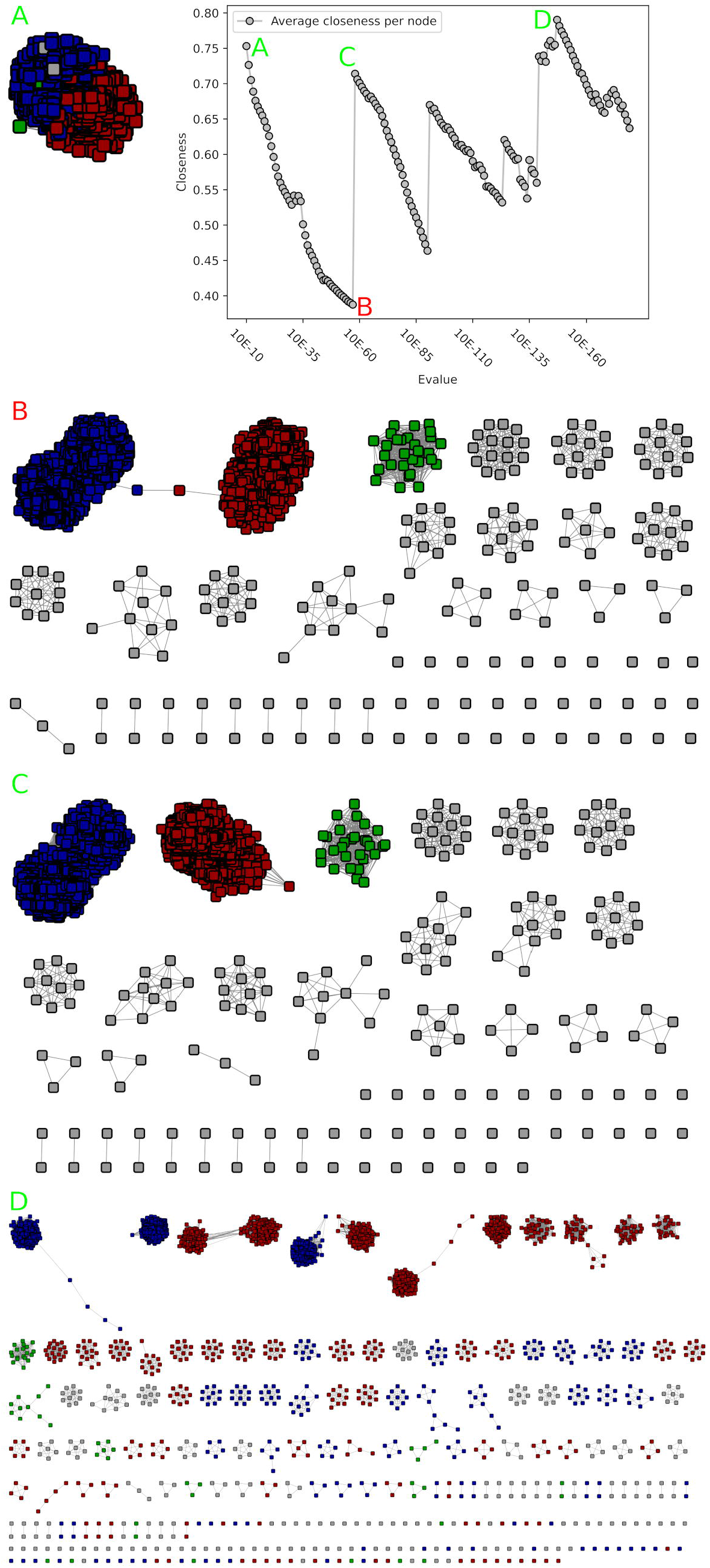
Closeness value and corresponding networks of the GH51 SSN analysis. As with GH68, multiple configurations could be of interest. A) The network is still only a single component. B) at 10^−57^, two main groups are sparsely connected, and the closeness value reaches its lowest point. C) At 10^−58^ these networks are disconnected, and the closeness value increases. D) 10^−147^ represents the highest peak after 10^−58^, and many more possible subfamilies are visible. All represented networks have been scaled down to 25% of their nodes, to make visualization possible. The network of A) was scaled down to 10%.

At 10^−91^, subfamily 1 and 3 split into two subfamilies, both containing characterized enzymes with similar activity/EC numbers. The split of subfamily 3 seems driven by the phylogeny, and resulted in 139 unclassified sequences as result of the split of subfamily 1. As a consequence, based on e-value range and functional specialization, 12 subfamilies of GH51 were build according to the 10^−58^ cutoff discussed above, including the three main subfamilies and nine small subfamilies among the 55 (having more than ten members; total 216 sequences) without functional evidence so far.

### Iterative SSNs might be necessary

The investigation of family GH45 initially resulted in 2467 modules sequences after fragment filtering. This only corresponded to a reduced fraction (77%) of the 3208 proteins annotated as GH45 and in-depth analysis revealed that the GH45 HMM did not allow to extract complete modules in many cases. Such situation might result from extreme sequence diversity within a family, for which a single model will necessarily be highly degenerated on the most variable regions. An initial SSN was thus built on the 2467 proteins that passed the filter, and the closeness criterion used to determine a first optimal split at 10^−7^ allowing the generation of two subgroups. Two HMMs were built (one for each subgroup) and used to successfully extract 3200 more complete modules (passing the filter) among the 3208 GH45 sequences. Therefore, after one iteration, a second SSN was built, and analyzed with the closeness criterion.

There are three known activities in family GH45, namely endoglucanase (EC 3.2.1.4), xyloglucan-specific endo-β-1,4-glucanase (EC 3.2.1.151) and endo-β-1,4-mannanase (EC 3.2.1.78). Notably 13 enzymes display dual 3.2.1.4/3.2.178 activities, one enzyme dual 3.2.1.4/3.2.1.151, and another enzyme dual 3.2.1.78+3.2.1.151. A previous analysis identified 3 subgroups for this family (16). Closeness centrality identified, after the initial 10^−6^ cutoff, another a single additional optimum at 10^−26^ (Figure 5). At this cutoff, four major groups (>100 sequences; one with approx. 2000, two of ∼440-510 and one of ∼240) were identified, corresponding to the previously identified three groups (16) and an additional group. This new group (∼440 members) consists nearly exclusively of Basidiomycetes sequences, which are globally more distant from other GH45 sequences, and have no characterized member yet. In the previous publication by (16) this group has not been seen, due to the small number of available Basidiomycetes sequences (only one sequence present in the previous analysis; XP_761686 in group C). This group did not appear in the first SSN, since the initial HMM did not detect a domain in all of its sequences. The sequences caught in the second SSN round were not limited to this group but also distributed over other small groups, therefore showing that the iterative SSN expanded multiple groups. No separation of EC numbers could be seen, with the largest group containing all three EC numbers, and the two smaller groups containing only the EC 3.2.1.4. Even at low e-values, e.g. 10^−111^, no full separation of the three activities could be obtained.

**Figure 5:**
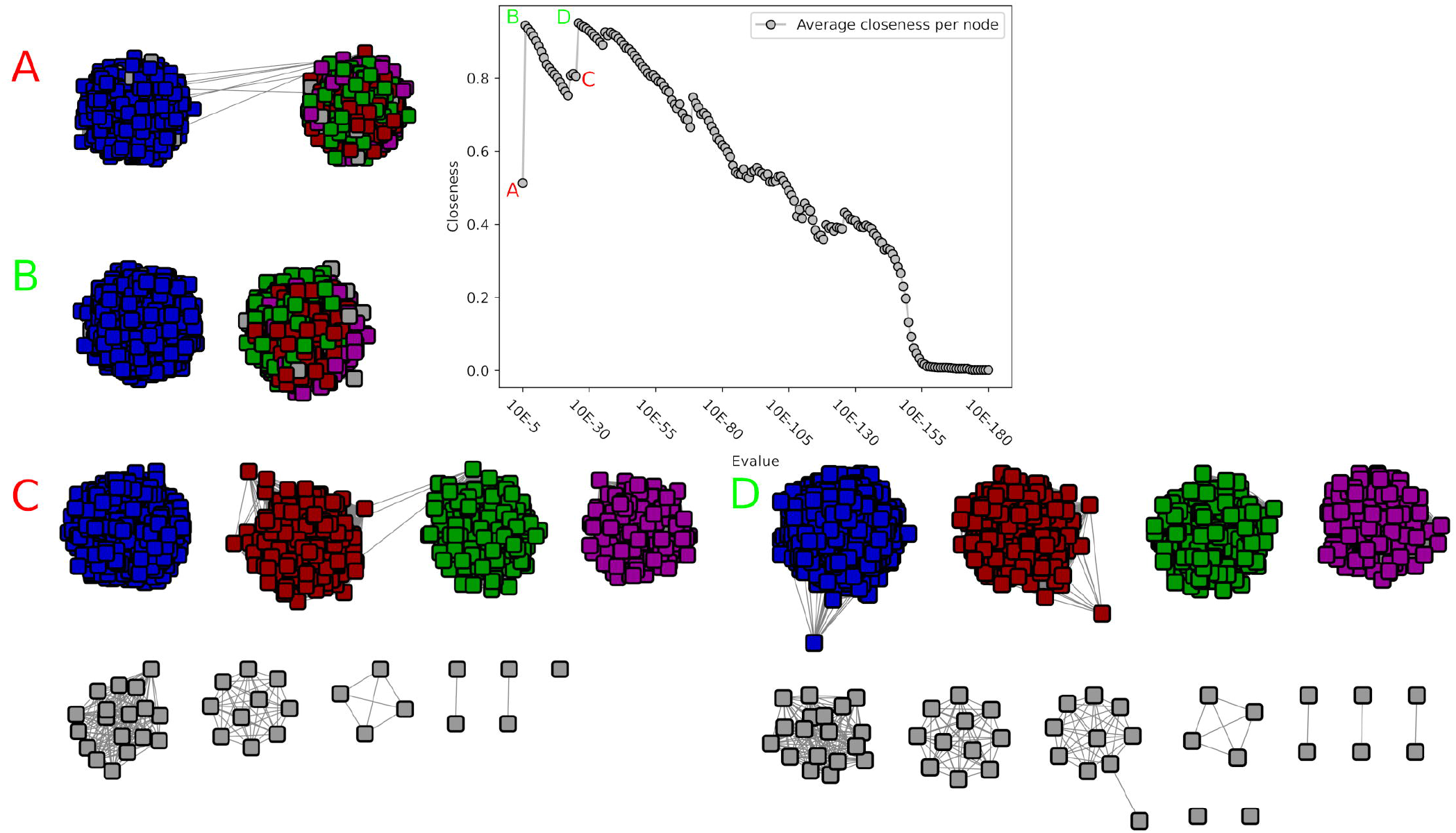
Closeness value and corresponding networks of the second GH45 SSN analysis. A) At 10^−5^ the network is still one component, although sparsely connected. B) At 10^−6^ the two subnetworks are disconnected and the closeness value increases. C) Another subnetwork has emerged, and two more are visible. D) At 10^−26^ the closeness value reaches its absolute maximum at 0.95 and four major subnetworks are visible.

### Functional versus structural separation into subfamilies

Family GH19 was investigated, following a recent publication about its putative subfamilies (17). We obtained 11,290 sequences after filtering, with in total two different activities, chitinase (EC 3.2.1.14) and lysozyme (EC 3.2.1.17) across 97 characterized enzymes. Closeness centrality identified multiple possible optima for subfamily separation (Figure 6). The first optimum at 10^−26^ (closeness of 0.57) splits the family into two main subgroups which display a single and specific activity, as well as 5 smaller subfamilies (>=10 sequences) and 0.9% unclassified sequences. Some lower e-value cutoffs lead to higher values of closeness centrality (10^−58^ reaches 0.58; 10^−76^ reaches the global optimum with 0.77) and would theoretically result in a better subfamily separation. It however leads to much more subfamilies (62 and 88 total with more than 10 members resp.), much more unclassified sequences (7% and 13% resp.), while no further separation of functions was observed, since known functions are already optimally split at 10^−26^. As discussed in (17), deeper subfamily splits result in a separation of structural features, and not of functions. We therefore decided to split GH19 into only two subfamilies.

**Figure 6:**
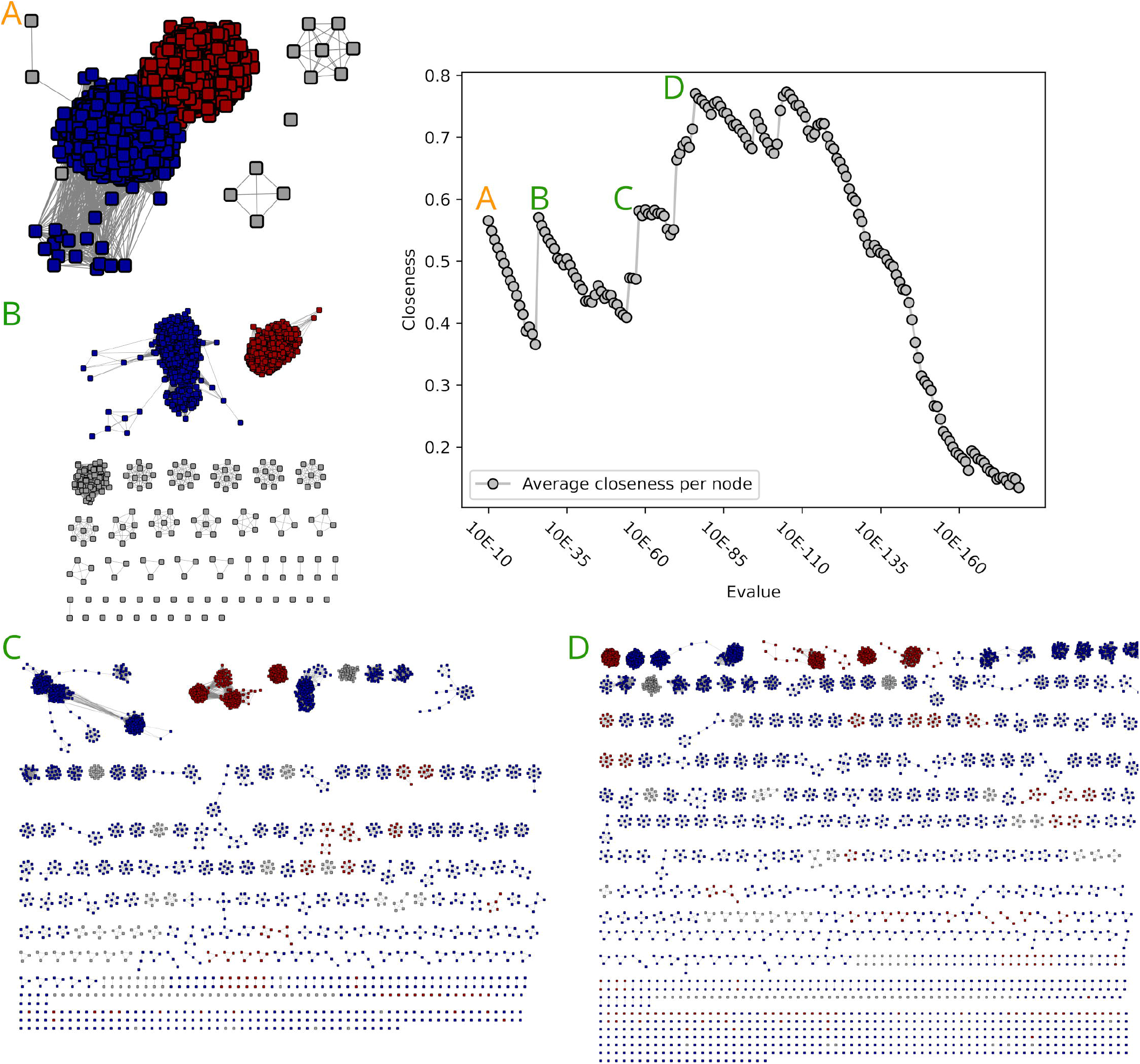
Closeness value and corresponding networks of the GH19 SSN analysis. As with GH68 And GH51, multiple configurations could be of interest. A) The network is still mostly a single component. B) At 10E-26 two main networks are visible. C+D) 10E-58 and 10E-76 are other optima, showing a vast amount of subnetworks. All networks have been scaled down to 10% for visualization purposes.

### Not all protein families have subfamilies

While we were able to detect subfamilies for several families, some did not seem to display subfamilies based on the available data, such as families PL26 and GH54. For PL26 we obtained 1657 sequences after filtering, with only one characterized enzyme (EC 4.2.2.24). The closeness centrality value started perfectly at 1 for an e-value threshold of 10^−5^ and decrease very slowly up to 0.95 before a local optimum (closeness back to 1) at 10^−46^, where one small group of 21 sequences emerged (Figure 7). This group did not contain any known activities, and consisted out of 20 sequences from the phylum Bacteroidetes and one of the phylum Balneolaetoa. At the highest e-value, 10^−179^, the PL26 network would still consist out of one main subnetwork of 1632 nodes, the small subnetwork of 21 nodes that appeared at 10^−46^, two linked sequences and two singles.

**Figure 7:**
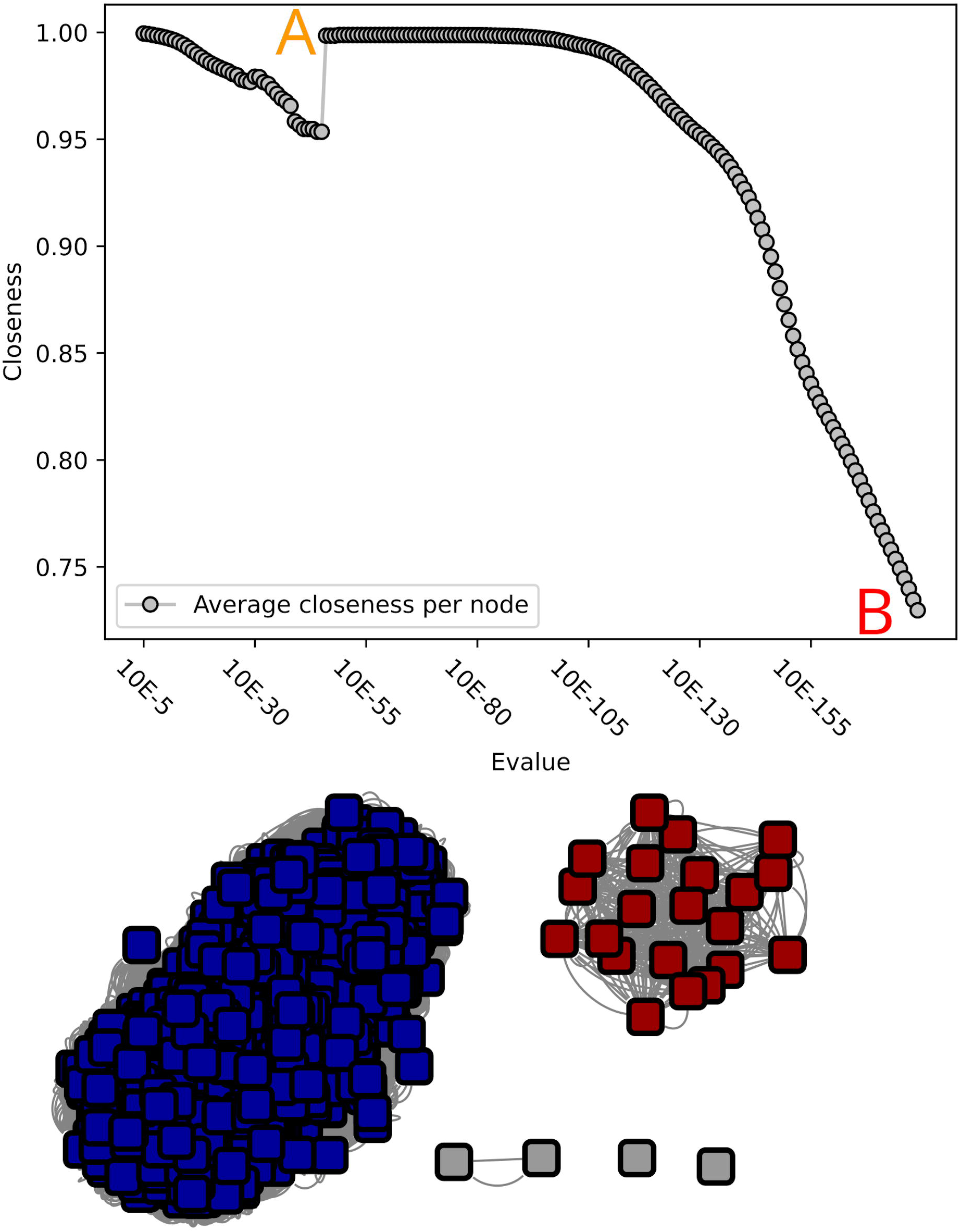
Closeness value and corresponding networks of the PL26 SSN analysis. A) The small red network is split off the main blue network B) Final network as depicted below the graph, at an evalue of 10^−179^.

For GH54 we obtained 1594 sequences after filtering, including 20 sequences with known activities (19 with α-L-arabinofuranosidase, EC 3.2.1.55; one with dual α-L-arabinofuranosidase and β-xylosidase, EC 3.2.1.37 activities). In this case, closeness centrality also starts at 1 for the highest e-value cutoffs, stays stable until 10^−67^ (closeness centrality of 0.99) and slowly decreases until 10^−131^ (closeness of 0.70) before raising back to a local optimum at 10^−139^ (closeness of 0.92; Figure 8). At this lowest cutoff, the largest subnetwork contains 1376 out of 1594 sequences, while the 216 remaining sequences spread over 25 smaller subnetworks (maximum size 86; only four with more than ten sequences). These subgroups follow taxonomic splits, and no functional division could be seen, since all characterized enzymes were contained in the main network.

**Figure 8:**
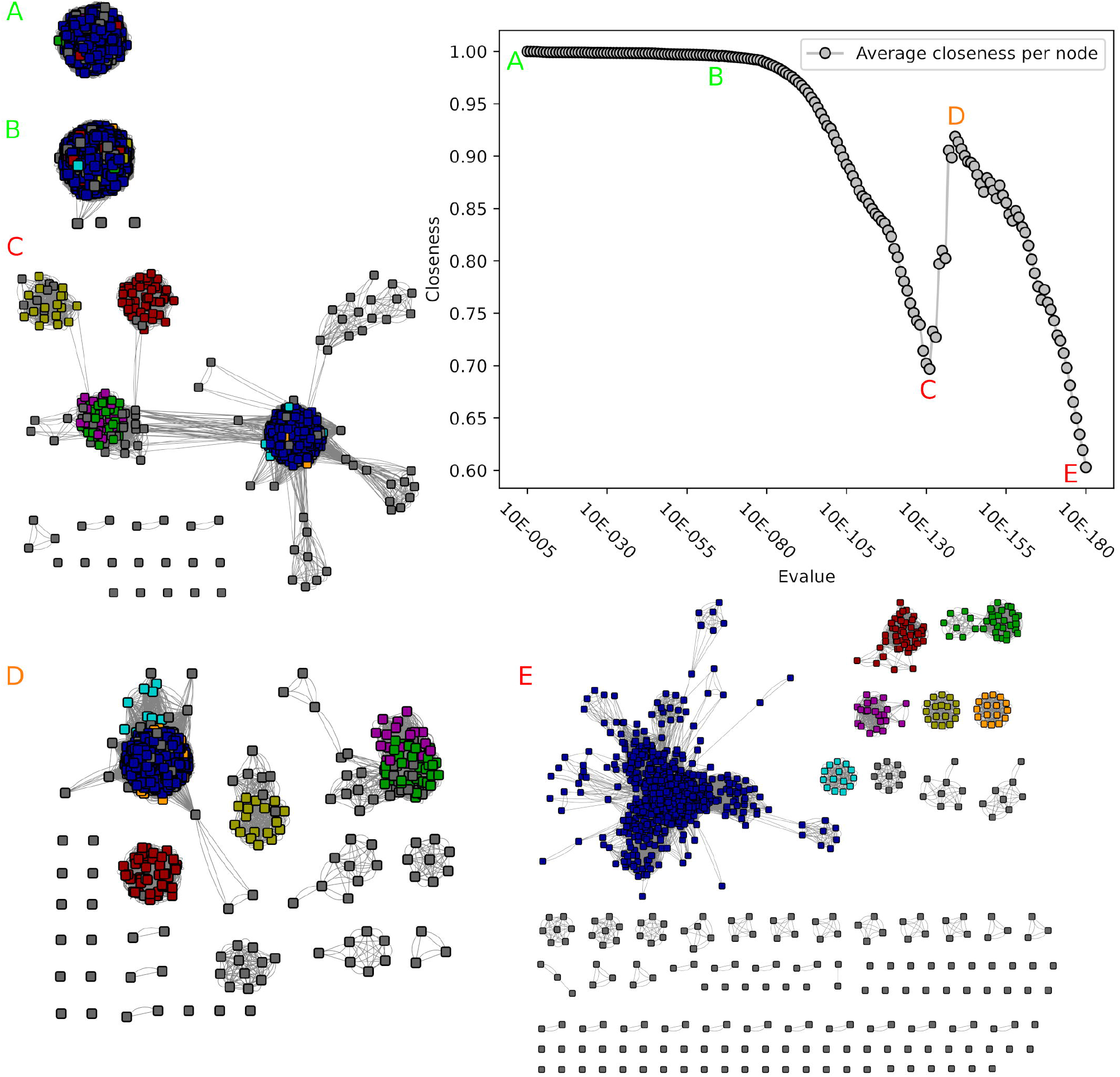
Closeness value and corresponding networks of the GH54 SSN analysis. A) The network is a single component, no subnetworks are visible. B) The closeness value changes slightly, and two disconnected nodes emerge. C) The closeness value reaches its first local minimum. While subgroups are visible, they are all still connected. D) The closeness value reaches its only peak. The main network still contains 86% of all sequences, and the smaller networks all follow phylogeny. E) At the maximum evalue of 10^−179^ various smaller networks emerge, yet no sensible bigger subnetworks are visible.

We therefore concluded that based on the available data, and given the absence of characterized members among emerging small groups, no subfamily should be created for both PL26 and GH54.

## Discussion

The goal of this work was to identify an objective and performant criterion to facilitate the exploration of SSNs by expert curators during an analysis of subfamily delineation. We demonstrated on artificial data that the *closeness* centrality criterion displays encouraging behavior, given that it favors perfect subnetworks, and penalizes sparsely connected networks, which is exactly what is desired for protein subfamilies translated into a SSN. We showed that the suggested optimal values are extremely close to those manually chosen by expert curators in previously published analyses, who sometimes identified several possible thresholds/subdivisions based on phylogenetic levels. In most cases, closeness centrality also identified several optima. In the analyzed families, optima of lowest e-values reflected the taxonomic separation of the sequences and likely strict orthologous groups, while more functionally-relevant splits were obtained at higher thresholds. This likely reflect the evolution of proteins with ancient duplication leading to neofunctionalization bringing fitness to various surviving species, while taxonomic groups are accentuated by species survival/extinction, and sometimes by a bias in genome sequencing. The existence of several optima still raises the question of the limitation of defining a single level of “subfamily”. A more general framework with flexible number of sub-, subsub-, or more levels of classification might be relevant to have. This would allow (i) a better coverage of the sequence space, as many sequences cannot be assigned to a subfamily at lower thresholds; and (ii) finer functional prediction, associated to the diversity of the functions in the subdivision, hence broader at higher thresholds. In some cases, this might also depend on the scope of the database. For the specialized GH19 database (17), the authors chose to annotate multiple levels of subfamily, corresponding to first functional and later structural differences. The CAZy database is primarily interested in the functional differences, and we therefore decided that one subfamily-level annotation is sufficient, although this might depend on the families or enzyme classes. Therefore, we only implement one subfamily hierarchy level, as do many other specialized databases like MEROPS (18), ESTHER (19), or SulfAtlas (20). The subfamily division is however not applicable to all families in a systematic/automatic manner, as it requires in-depth analysis to ensure long-term stability of the proposed classification, the availability of enough characterized members and human decision when several optima are found. This highlights the need to support such expert databases, which are dedicated to mine the functional annotation of (meta)genomes in the literature, as such effort can no longer be supported by universal databases.

For GH55, we found an agreement to prior analysis (15). Bianchetti et al. Identified, in a large-scale screening of GH55, three major subfamilies. Two of these families contained functional enzymes, and were divided into a fungal and a bacterial family, with the third family containing non-functional bacterial proteins. These three families correspond to the three groups identified by our SSN investigation, although we did not further subdivide based on phylogeny, as Bianchetti et al. did (15). In this case we can not only show that SSNs are able to delineate functionally different enzymes, in some cases they also might be able to delineate non-functional enzymes, as seen by the identification of a subfamily where no functions could be found even after testing. This potentially allows to save time in future endeavors by ruling out known non-functional proteins.

Our analyses also globally agreed with prior investigations of families GH68 (21), GH51 (22) and GH45 (16). In case of GH68, this family would separate into two subfamilies and does not result into any unclassified sequences. A possible further in-depth classification, as also suggested by Velazquez-Hernandez, could be considered in the future, in case new functions are discovered in this more distant and small groups.

For GH51, three main subfamilies, corresponding to the three main clades identified in (22), will be created, as well as nine smaller ones. The two biggest subfamilies, representing 96% of the data, are distinguished by a structural feature in the sequences. One protein in one of the subfamilies contains a minor side function. Further biochemical characterization is necessary to see if this is a property of the whole subfamily and if it might be a distinguishing factor between the both major subfamilies. The third major subfamily represents a distinct function, whereas the fourth is of unknown function, but is characterized by the presence of a carbohydrate binding domain, indicating considerable differences within this family.

The GH45 case differs from the previous cases as none of the subfamilies demonstrated separate biochemical functions. Yet four subfamilies will be implemented, since this family seems to be very diverse, and the initial HMM for the detection of GH45 did not allow a full detection of module sequences. The subfamily HMMs yield an improved performance and better detection, making their implementation therefore necessary.

For some families, we were not able to identify any sensible subgrouping. We hypothesized that two or more groups could exist in GH54, because most enzymes in GH54 contained only one function, but one enzyme from *Trichoderma koningii* has been deemed to have an dual activity (additional β-xylosidase activity) (23). The SSN analysis indicated that no such major subgroups exist though. While reviewing the literature, it appears that this β-xylosidase activity is only a minor side-activity, which also was reported for other enzymes in this group before (24,25). We therefore conclude that no subgroups exist in GH54, and that potentially all members might have a dual activity.

Family PL26 was included in our analysis, since it is the largest PL family, for which no subfamilies were previously created (26). After the creation of the SSN, we were unable to detect any major subfamilies. Given that only one protein has been characterized so far, it might be that this family truly does not contain any more functions or subfamilies.

A limitation of the *closeness centrality* is that it is independent of the amount of subnetworks and of their size. Two perfect subnetworks of the size 50 will give the same outcome as 50 perfect subnetworks of size 2. An option would be to penalize subnetworks below a certain size, typically below the required family size, by downgrading their values e.g. assigning a value of 0 as for singlets. While this might be necessary in case many different networks are built at the same time, in most circumstances a manual evaluation by the curator with the amount of networks in mind should suffice.

An unexpected limitation which we encountered is the usage of the e-value for the generation of SSNs. Most researchers decrement the e-value to generate their SSNs (4,5). The e-value is a floating point value, where reaching 0.0 is considered to be the most significant value, indicating a very good match, most often resulting in perfect matches. The e-value is dependent on database size and query length though. In case of too large families, or very long protein domains (e.g. PL26s average is 867 AA), the e-value will often result in 0 (see supplementary Figure S3, with an overview of e-values per percentage identity), due to the limit of 10^−308^ for a 64-bit floating point value with double precision (IEEE 754 standard for floating point arithmetic, (27) ; the equivalent limit for 32-bit is 10^−38^). Distribution of e-values can often be seen with a maximum of 10^−160^ or 10^−180^, with more significant values being 0 for long domains. For these long domains this can lead to many hits, which identity percent can reach fairly low/limit values between 20-30%, and yet lead to an e-value of 0. In case of PL26, e-values of 0 were obtained at worst with a sequence identity of 32%. Bitscores (or a combination of identity percent and coverage control), should rather be used instead of e-values in case of unusually long sequences.

We also tried to evaluate a proposed mechanism for network filtering from the field of neuroscience (28), but this proved to be computationally infeasible, with runtimes of more than 3 days for comparatively small networks from our collection. We furthermore hoped to aid our annotation efforts with well-known algorithms for grouping data points, like clustering or PCA, despite them not being favored in earlier research (29). It turned out that even for rather simple group combinations with rather little sequences, like two or four groups in GH68, these algorithms are already severely challenged (see supplementary results). At this point it needs to be evaluated if the results from the other techniques like PCA are correct, and the results based

on *closeness centrality* might be incorrect. A manual inspection with Cytoscape (30) proofed to be efficient and simple for the small SSNs, and confirmed that the results obtained by the evaluation of *closeness centrality* are indeed correct. Overall, these results from the other techniques make our exploration of other criteria even more worthwhile. We hope that the *closeness centrality* metric will be used during other research projects in this field, to combine expert knowledge with reproducible technical measures.

## Materials and Methods

### Generation of artificial data

Artificial data was generated with the Python networkX 2.4/5 library (31).

We first used the network generators to create different *subnetworks* (to be connected during the second step): (i) two perfect networks (all possible edges between all nodes exist) of size 50 and 200, using the *nx*.*complete_graph* function; six random networks of size 200 with 300, 500, 1000, 2000, 3000 and 4000 edges (representing from 1.5% to 20.1% of a complete graph) using the *nx*.*gnm_random_graph* function; and two networks consisting out of 50 or 200 single nodes, or singletons.

Secondly, we generated network *collections*. A *collection* consists in multiple *networks* build upon the same initial pair of *subnetworks* (described above; see supplementary table 2). A *collection* starts with a first *network* simply consisting in the two disconnected *subnetworks*. The collection is completed by additional *networks* obtained by randomly connecting the nodes of one *subnetwork* to the other with an increasing number of edges, from one to the maximum possible (see Supplementary Figure S1A). This means that a combination between a *subnetwork* of size 50 and another of size 200 would result in a *collection* that contains 10,001 *networks*, one without connection and then from a single connection between a random node of each *subnetwork*, up to the connection all 50 nodes of the first *subnetwork* to the 200 nodes of the second *subnetwork*. In case of two perfect *subnetworks*, the lastly created *network* in the *collection* would also be a perfect network.

In the specific case of a single-node *subnetwork* combined to a random or perfect *subnetwork*, two additional strategies of connections were performed to better explore the space of topologies, while ending in the same number of *networks* in the *collection*: 1) all single nodes (initially one and lately all) were first randomly connected to nodes of the other *subnetwork*. Once all connected, a second edge was added to each of these nodes, and this was repeated until all connections were made (see Supplementary Figure S1B); 2) a random node of the single-node *subnetwork* was first connected to all nodes the other *subnetwork* (initially to one and lately to all), and only afterwards a second single node was connected in the same way, until all connections were made (see Supplementary Figure S1C).

Not all possible *subnetwork* pairs were used to generate *networks/collections*, as only the following cases were considered: perfect network of size 50 randomly connected to all possible *subnetworks* (single node of size 1, 50 and 200; perfect subnetworks of size 3, 50 and 200, random networks of size 50 with 250 and 500 edges, and random networks of size 200 with 500 to 4000 edges) and single node networks (size 50 and 200) connected to random networks of size 200 with 500 to 4000 edges with the three connection strategies. An overview is given in supplementary table 2.

### Network properties

Before the execution of any computation, the networks were converted to an undirected graph. Afterwards, various network properties were calculated, using networkX, and investigated for local and global minima and maxima. These 19 properties include the number of components, edges, s-metric, average degree centrality, average closeness centrality (with the parameter *wf_improved* -allowing to cope with disconnected components -either True and False), average betweenness centrality, average degree, average clustering, eigenvector centrality, betweenness centrality source, edge betweenness centrality, load centrality, edge load centrality, subgraph centrality, harmonic centrality, global reaching centrality, the amount of local bridges, the shortest path and asyn_lpa_communities (32,33). For the *artificial* data, all these properties were calculated, while CAZy families GH16 (8), GH19 (17), GH45, GH51, GH54, GH55, GH68, GH130 (14) and PL26, as well as different families of the Structure Function Linkage Database (SFLD) (3), SFLD families 19, 122 and 159 and Interpro IPR037434, were investigated with the closeness criterion only (see results).

A script for easier accessibility is available under https://github.com/bastian-wur/closeness_by_component.

### Generation of SSNs

Sequence similarities on SFLD and CAZy families were computed with Blastp v2.5 (34). Standard parameters were used, besides the parameter *max_target_seqs*, which was scaled to the size of the family/target database. The Blastp pairwise hits, based on e-value cutoff, were transformed into a digraph with the python library networkX v2.4/2.5 (31), while sequences without any hit were added as single nodes. We generated SSNs for each e-value cutoff, from 10^−5^ up to 10^−160^, by step of 10^−1^. Exceptions to these start-and end-points were made when either the network did not form any more subnetworks at high cutoffs, or when the network disintegrated into too many subnetworks (typically >1000) at low cutoffs.

All networks were visualized with Cytoscape version 3.7.2 (30). Graphs showing the evolution of the network properties were generated with Matplotlib (35-37).

GNU parallel version 20161222 has been used during this and other steps of this study (38).

### EFI-EST

The EFI-EST website (39) was used to reproduce the published IPR037434 family analysis, similarly to (10). An initial e-value cutoff of 10^−5^ was used, and subnetworks were produced based on this network by decreasing the e-value cutoff by steps of 10-5 until the maximum selectable e-value of 10^−100^. Steps of 10^−1^ were prohibitive in an online-tool without the possibility of automation, and edge values were not included into EFI-EST SSNs during our first tests.

For reading xgmml files, networkxgmml v0.1.6. was used (https://pypi.org/project/networkxgmml/).

### Extraction of domain sequences

SFLD sequence data for families 19 of glutathione transferase (12), family 159 of tautomerase (11), and family 122 of nitroreductase (13) were downloaded from http://sfld.rbvi.ucsf.edu/archive/django/superfamily/index.html (9). For these families the full protein sequences were used.

Data for glycoside hydrolase for families GH16, GH19, GH51, GH54, GH55, GH68, GH130 and polysaccharide lyase family PL26 were directly extracted from the CAZy database (40), and were further supplemented with data by collaboration partners. Domain sequences were extracted from the full proteins using in-house pipeline based on the results of *hmmscan* function in HMMer3.3 (41) as follows. HMM searches using the in-house CAZy HMM profiles were performed and only matches with an e-value <1-4 were further processed. The sequence and HMM coordinates were extracted in an attempt to reconstruct a domain sequence from fragments created by the local detection of HMM searches. Two consecutive fragments were assembled only in the given conditions: (i) sequence start coordinates must be separated by >20 amino-acids, as well as sequence end coordinates; (ii) sequence end of the first fragment should not be separated from the start of the second fragment by >200 amino acids; (iii) HMM coordinates must not overlap (end of the first vs start of the second) by more than 30 amino acids. Any domain sequence, initially selected or reassembled, shorter than half of the HMM length was discarded. A manual inspection was performed for sequences longer or shorter than the average sequence length +/-three standard deviations, to discard gene models issues. In all cases >95% of the original input sequences passed the screening, except for GH45 (see Results section). For the construction of the second SSN for GH45, domains were extracted as described but using two GH45 HMMs. In case both HMMs identified a domain in the same protein, the longer domain was used.

### Minimized version for visualization

Since many of the generated networks were too large to be visualized in Cytoscape, and preliminary clustering seemed to warp network structure in some cases, networks for visualization were generated differently. A custom and modified version of uniform edge sampling with graph induction was implemented (42), to generate representative smaller graphs from the full graph.

To retain network structure, subgraphs were generated per connected component. Components smaller than a chosen node amount (in most cases 10) were retained as they were. Larger components were sampled down to a specified percentage (in most cases 25 or 50%; exact values specified per figure), with respect to the minimum amount of nodes of small components. The subsampled graphs, initially empty, were completed by random edges (Python *random*.*choice* function) and their nodes sequentially until the minimum amount of nodes was reached. At last, all edges connecting the sampled nodes in the initial component were added as well (42). In case a component was split by the random subsampling, reconnection was realized as follows. Iterating over all split sub-components, the shortest path algorithm (as implemented in networkX) was used to determine the shortest path between these selected nodes based on the initial component, and all necessary edges were iteratively added. The computation was stopped when the component was again fully connected.

## Supporting information

Supplementary methods and results

## Data availability

New subfamilies are published on the CAZy website, www.cazy.org.

## Acknowledgments

The authors would like to thank Vincent Lombard for technical support and for administration of the CAZy cluster. The authors would furthermore like to thank Bernard Henrissat for proofreading and providing useful comments.

## Author contributions

Conceptualization: BVHH

Data curation: BVHH, NT

Formal analysis: BVHH

Funding acquisition: NT

Investigation: BVHH

Methodology: BVHH, NT

Project administration: NT

Resources: NT

Software: BVHH

Supervision: NT

Validation: BVHH

Visualization: BVHH

Writing – Original draft preparation: BVHH, NT

Writing-Review and editing: BVHH, NT

## References

1. Schnoes AM, Ream DC, Thorman AW, Babbitt PC, Friedberg I. Biases in the Experimental Annotations of Protein Function and Their Effect on Our Understanding of Protein Function Space.. PLoS Comput Biol. 2013 May 30;9(5):e1003063.

2. Jiang Y, Oron TR, Clark WT, Bankapur AR, D’Andrea D, Lepore R, et al. An expanded evaluation of protein function prediction methods shows an improvement in accuracy. Genome Biol. 2016 Dec;17(1):184.

3. Holliday GL, Brown SD, Akiva E, Mischel D, Hicks MA, Morris JH, et al. Biocuration in the structure–function linkage database: the anatomy of a superfamily. Database. 2017 Jan 1:bax006

4. Brown SD, Babbitt PC. New Insights about Enzyme Evolution from Large Scale Studies of Sequence and Structure Relationships. J Biol Chem. 2014 Oct;289(44):30221–8.

5. Gerlt JA, Bouvier JT, Davidson DB, Imker HJ, Sadkhin B, Slater DR, et al. Enzyme Function Initiative-Enzyme Similarity Tool (EFI-EST): A web tool for generating protein sequence similarity networks. Biochim Biophys Acta. 2015 Aug;1854(8):1019–37.

6. Fetrow JS, Babbitt PC. New computational approaches to understanding molecular protein function. PLoS Comput Biol. 2018 Apr 5;14(4):e1005756.

7. Drula E, Garron M-L, Dogan S, Lombard V, Henrissat B, Terrapon N. The carbohydrate-active enzyme database: functions and literature. Nucl Acids Res. 2022 Jan 7;50(D1):D571–7.

8. Viborg AH, Terrapon N, Lombard V, Michel G, Czjzek M, Henrissat B, et al. A subfamily roadmap of the evolutionarily diverse glycoside hydrolase family 16 (GH16). J Biol Chem. 2019 Nov;294(44):15973–86.

9. Akiva E, Brown S, Almonacid DE, Barber AE, Custer AF, Hicks MA, et al. The Structure–Function Linkage Database. Nucl Acids Res. 2014 Jan;42(D1):D521–30.

10. Travis S, Shay MR, Manabe S, Gilbert NC, Frantom PA, Thompson MK. Characterization of the genomically encoded fosfomycin resistance enzyme from Mycobacterium abscessus. Med Chem Commun. 2019;10(11):1948–57.

11. Mashiyama ST, Malabanan MM, Akiva E, Bhosle R, Branch MC, Hillerich B, et al. Large-Scale Determination of Sequence, Structure, and Function Relationships in Cytosolic Glutathione Transferases across the Biosphere. PLoS Biol. 2014 Apr 22;12(4):e1001843.

12. Davidson R, Baas B-J, Akiva E, Holliday GL, Polacco BJ, LeVieux JA, et al. A global view of structure–function relationships in the tautomerase superfamily. J Biol Chem. 2018 Feb;293(7):2342–57.

13. Akiva E, Copp JN, Tokuriki N, Babbitt PC. Evolutionary and molecular foundations of multiple contemporary functions of the nitroreductase superfamily. Proc Natl Acad Sci USA. 2017 Nov 7;114(45):E9549–58.

14. Li A, Laville E, Tarquis L, Lombard V, Ropartz D, Terrapon N, et al. Analysis of the diversity of the glycoside hydrolase family 130 in mammal gut microbiomes reveals a novel mannoside-phosphorylase function. Microb Genom. 2020 Oct 1;6(10):mgen000404.

15. Bianchetti CM, Takasuka TE, Deutsch S, Udell HS, Yik EJ, Bergeman LF, et al. Active Site and Laminarin Binding in Glycoside Hydrolase Family 55. J Biol Chem. 2015 May;290(19):11819–32.

16. Igarashi K, Ishida T, Hori C, Samejima M. Characterization of an Endoglucanase Belonging to a New Subfamily of Glycoside Hydrolase Family 45 of the Basidiomycete Phanerochaete chrysosporium. Appl Environ Microbiol. 2008 Sep 15;74(18):5628–34.

17. Orlando M, Buchholz PCF, Lotti M, Pleiss J. The GH19 Engineering Database: Sequence diversity, substrate scope, and evolution in glycoside hydrolase family 19. PLoS ONE. 2021 Oct 26;16(10):e0256817.

18. Rawlings ND, Waller M, Barrett AJ, Bateman A. MEROPS : the database of proteolytic enzymes, their substrates and inhibitors. Nucl Acids Res. 2014 Jan;42(D1):D503–9.

19. Lenfant N, Hotelier T, Velluet E, Bourne Y, Marchot P, Chatonnet A. ESTHER, the database of the α/β-hydrolase fold superfamily of proteins: tools to explore diversity of functions. Nucl Acids Res. 2012 Nov 26;41(D1):D423–9.

20. Barbeyron T, Brillet-Guéguen L, Carré W, Carrière C, Caron C, Czjzek M, et al. Matching the Diversity of Sulfated Biomolecules: Creation of a Classification Database for Sulfatases Reflecting Their Substrate Specificity. PLoS ONE. 2016 Oct 17;11(10):e0164846.

21. Velázquez-Hernández ML, Baizabal-Aguirre VM, Bravo-Patiño A, Cajero-Juárez M, Chávez-Moctezuma MP, Valdez-Alarcón JJ. Microbial fructosyltransferases and the role of fructans. J Appl Microbiol. 2009 Jun;106(6):1763–78.

22. dos Santos CR, de Giuseppe PO, de Souza FHM, Zanphorlin LM, Domingues MN, Pirolla RAS, et al. The mechanism by which a distinguishing arabinofuranosidase can cope with internal di-substitutions in arabinoxylans. Biotechnol Biofuels. 2018 Dec;11(1):223.

23. Wan C-F, Chen W-H, Chen C-T, Chang MD-T, Lo L-C, Li Y-K. Mutagenesis and mechanistic study of a glycoside hydrolase family 54 α-L -arabinofuranosidase from Trichoderma koningii. Biochem J. 2007 Jan 15;401(2):551–8.

24. Guais O, Tourrasse O, Dourdoigne M, Parrou JL, Francois JM. Characterization of the family GH54 α-l-arabinofuranosidases in Penicillium funiculosum, including a novel protein bearing a cellulose-binding domain. Appl Microbiol Biotechnol. 2010 Jul;87(3):1007–21.

25. Saha BC, Bothast RJ. Purification and Characterization of a Novel Thermostable ⎵-L-Arabinofuranosidase from a Color-Variant Strain of Aureobasidium pullulans. Appl Environ Microbiol. 1998;64(5):216–220

26. Lombard V, Bernard T, Rancurel C, Brumer H, Coutinho PM, Henrissat B. A hierarchical classification of polysaccharide lyases for glycogenomics. Biochem J. 2010 Dec 15;432(3):437–44.

27. Muller J-M, Brisebarre N, de Dinechin F, Jeannerod C-P, Lefèvre V, Melquiond G, et al. Handbook of Floating-Point Arithmetic. Boston: Birkhäuser; 2010

28. De Vico Fallani F, Latora V, Chavez M. A Topological Criterion for Filtering Information in Complex Brain Networks. PLoS Comput Biol. 2017 Jan 11;13(1):e1005305.

29. Apeltsin L, Morris JH, Babbitt PC, Ferrin TE. Improving the quality of protein similarity network clustering algorithms using the network edge weight distribution. Bioinformatics. 2011 Feb 1;27(3):326–33.

30. Shannon P, Markiel A, Ozier O, Baglia NS, Wang JT, Amin N, et al. Cytoscape: A Software Environment for Integrated Models of Biomolecular Interaction Networks. Genome Res. 2003 Nov 1;13(11):2498–504.

31. Hagberg AA, Schult DA, Swart PJ. Exploring Network Structure, Dynamics, and Function using NetworkX. In: Proceedings of the 7th Python in Science Conference (SciPy 2008). 2008. p. 5.

32. Koschützki D, Schreiber F. Centrality Analysis Methods for Biological Networks and Their Application to Gene Regulatory Networks. Gene Regul Syst Bio. 2008;2:193–201.

33. Gómez S. Centrality in Networks: Finding the Most Important Nodes. In: Business and Consumer Analytics: New Ideas. Cham: Springer International Publishing; 2019.

34. Altschul SF, Gish W, Miller W, Myers EW, Lipman DJ. Basic Local Alignment Search Tool. J Mol Biol. 1990;215:403–10.

35. Harris CR, Millman KJ, van der Walt SJ, Gommers R, Virtanen P, Cournapeau D, et al. Array programming with NumPy. Nature. 2020 Sep 17;585(7825):357–62.

36. SciPy 1.0 Contributors, Virtanen P, Gommers R, Oliphant TE, Haberland M, Reddy T, et al. SciPy 1.0: fundamental algorithms for scientific computing in Python. Nat Methods. 2020 Mar;17(3):261–72.

37. J. D. Hunter. Matplotlib: A 2D Graphics Environment. Computing in Science & Engineering. 2007 Jun;9(3):90–5.

38. Tange O. GNU Parallel: The Command-Line Power Tool. ;login. 36(1):42–7.

39. Zallot R, Oberg N, Gerlt JA. The EFI Web Resource for Genomic Enzymology Tools: Leveraging Protein, Genome, and Metagenome Databases to Discover Novel Enzymes and Metabolic Pathways. Biochemistry. 2019 Oct 15;58(41):4169–82.

40. Lombard V, Golaconda Ramulu H, Drula E, Coutinho PM, Henrissat B. The carbohydrate-active enzymes database (CAZy) in 2013. Nucl Acids Res. 2014 Jan;42(D1):D490–5.

41. Eddy SR. Accelerated Profile HMM Searches. PLoS Comput Biol. 2011 Oct 20;7(10):e1002195.

42. Ahmed NK, Neville J, Kompella R. Network Sampling: From Static to Streaming Graphs. ACM Transactions on Knowledge Discovery from Data. 2014 8(2):1–56

