## Supplementary figures and images for "An objective criterion to evaluate sequence-similarity networks helps in dividing the protein family sequence space"

### Supplementary Figure 1

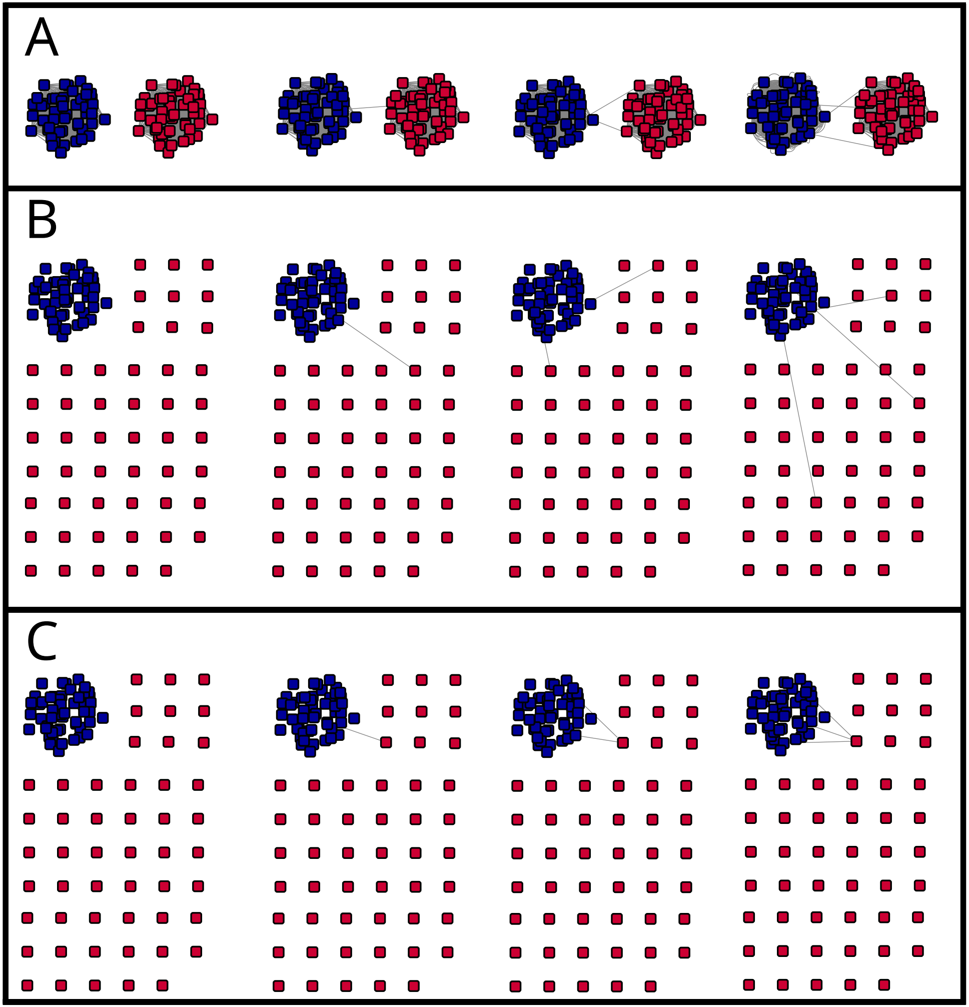

### Supplementary Figure 2

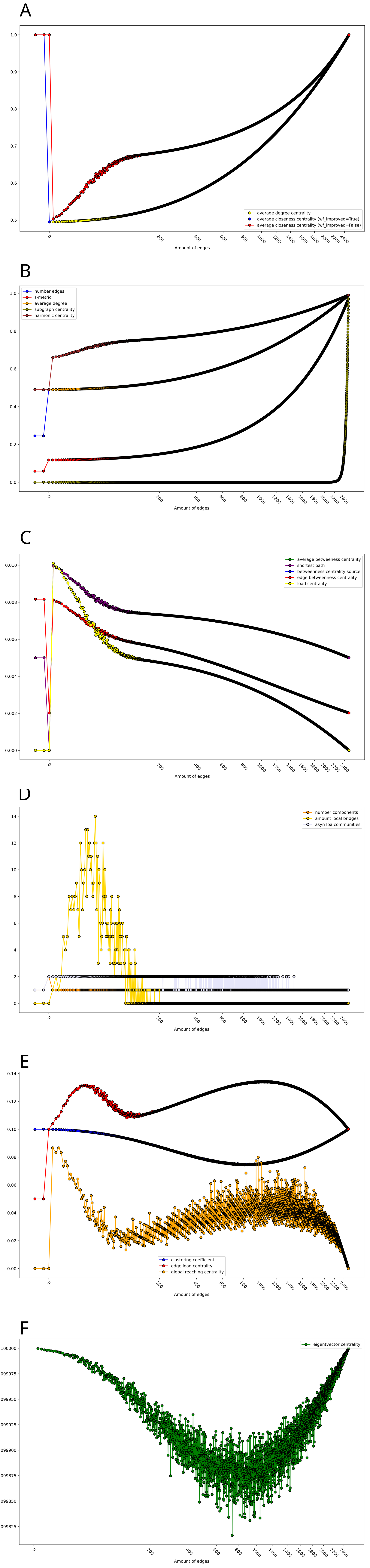

### Supplementary Figure 3

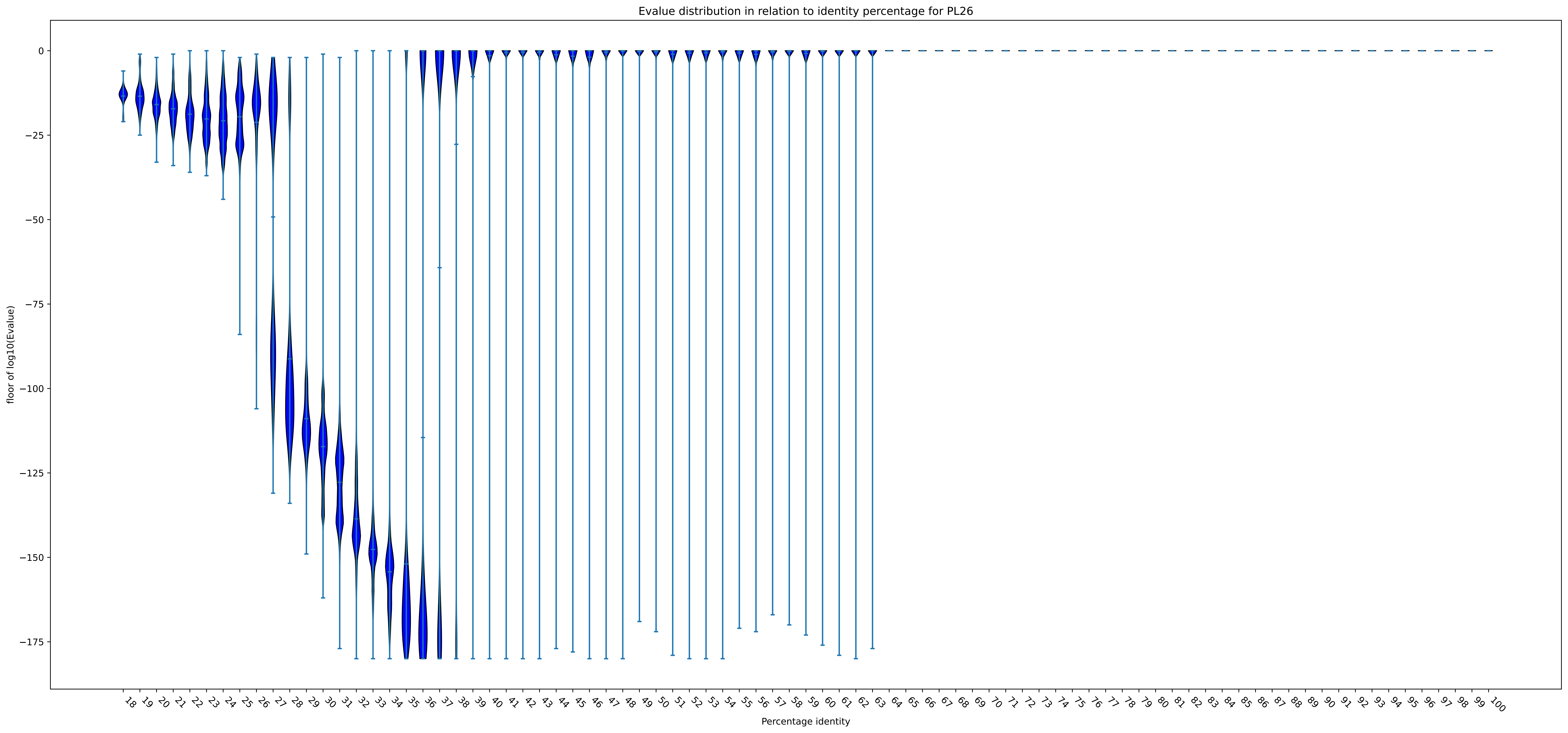

### Supplementary Figure 4

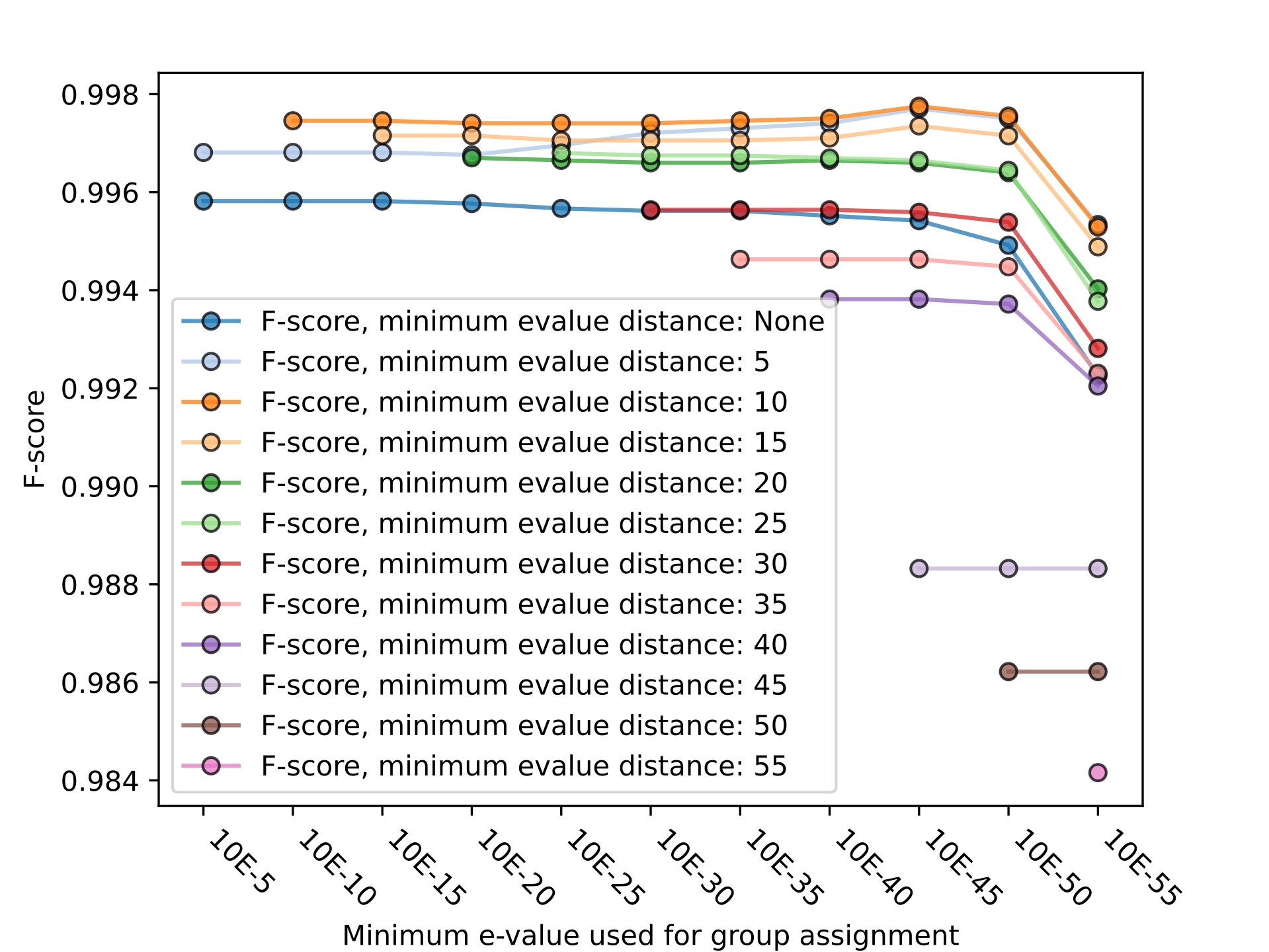

### Supplementary Figure 5

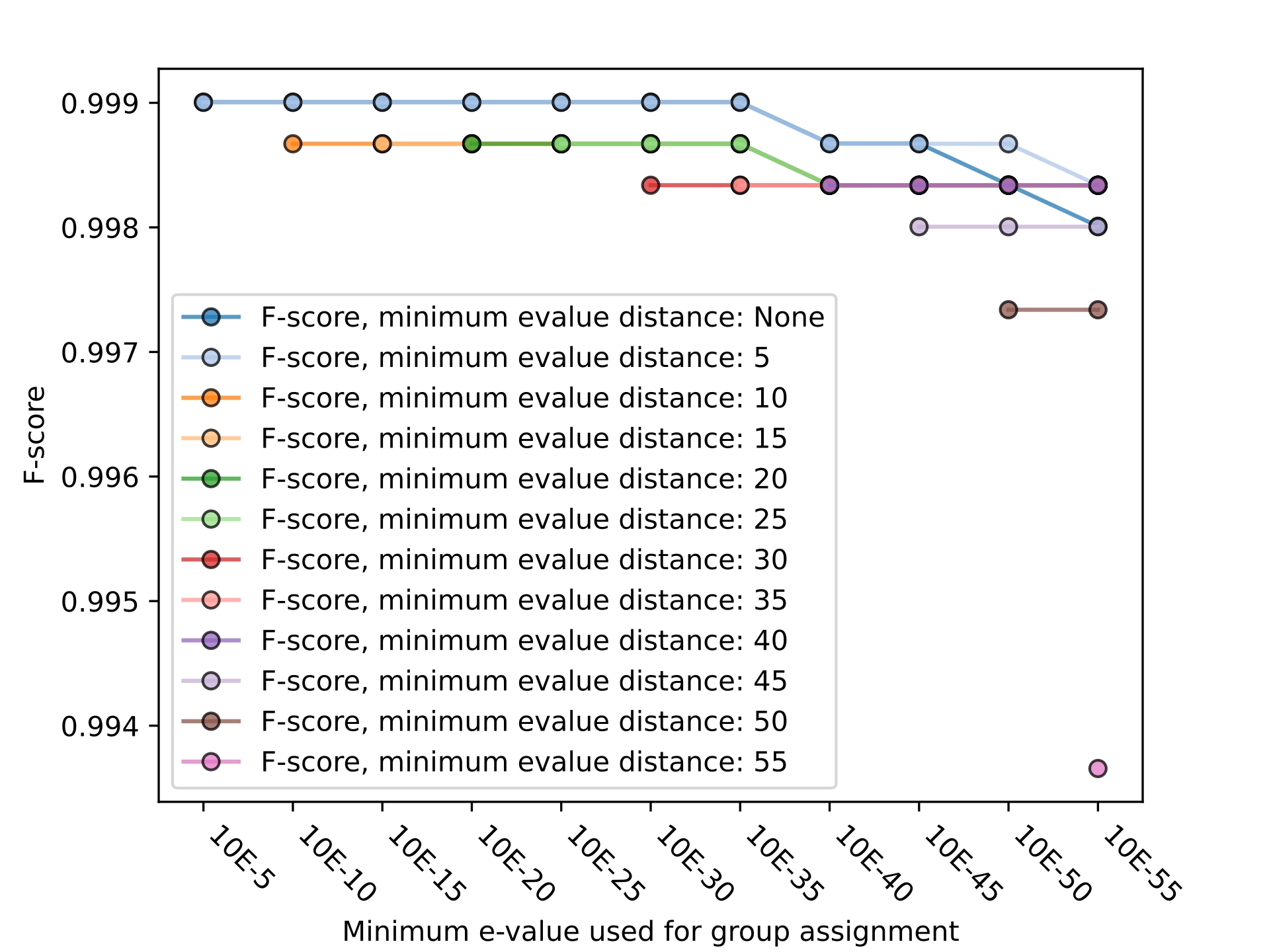

### Supplementary Figure 6

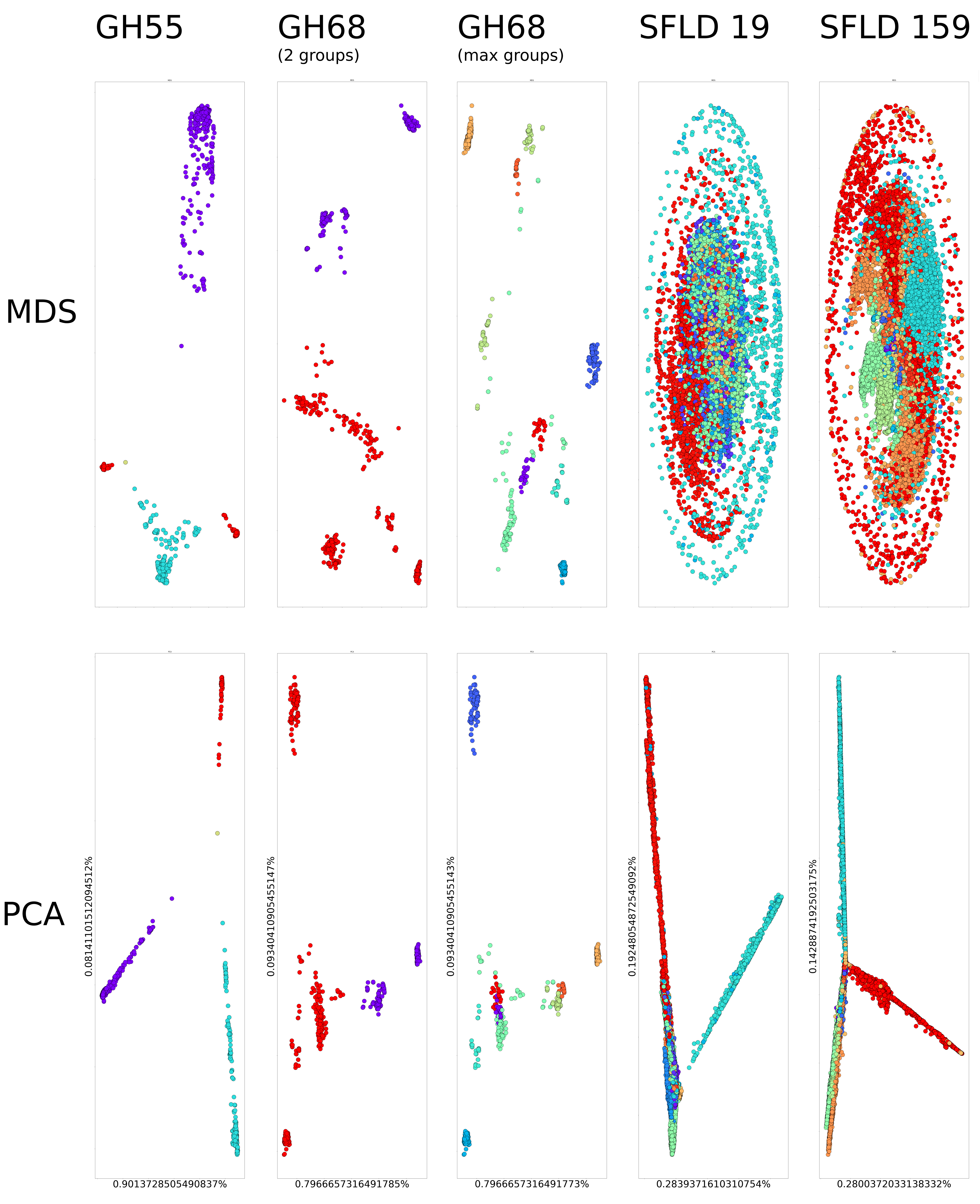
