## Supplementary methods and results for "An objective criterion to evaluate sequence-similarity networks helps in dividing the protein family sequence space"

B. V. H. Hornung[1,2,*], N. Terrapon[1,2,*]

[1] Aix Marseille Univ, CNRS, UMR7257 AFMB, Marseille, France

[2] INRAE, USC1408 AFMB, Marseille, France

* Corresponding author

Email: bastian.hornung @ gmx.de, nicolas.terrapon @ univ-amu.fr

### Supplementary Materials and Methods

#### Generation of subfamily HMMs

Based on selected closeness optima, HMM libraries were built and used for evaluation, similarly to the method described in Viborg et al. (1). For each SSN e-value threshold, connected components with >10 nodes were considered as putative subfamilies. A multiple sequence alignment (MSA) of each subfamily was built with Clustal Omega 1.2.4 (2) with standard parameters. A custom script was used to discard any columns in the MSA which contained more than 50% gaps. The resulting MSAs was used to build HMMs with HMMer (with standard parameters) and all subfamily HMMs assembled into an HMM library corresponding to the SSN threshold. An HMM search was thus performed over the full proteins, and each sequence was associated with the best scoring subfamily if satisfying two criteria: a minimal e-value and a minimal difference in e-value to the second best-scoring subfamily. Several annotation schemes were considered, using a minimal e-value from 0 to $10^{-55}$, and a minimal difference from 0 to $10^{-55}$. For each annotation schemes, the predicted subfamilies were compared to the SSN reference subfamilies, producing a confusion matrix and ROC curve by varying the minimal e-value cutoff (from 0 to $10^{-55}$).

#### Clustering

Clustering was performed with ELKI v. 0.7.6 (3). As input an asymmetrical matrix was used, which contained the absolute exponents of the evalues between the respective proteins. Values of 1 or 10 were equated with 0 or -1 respectively, if not match was found the value -2 was used. Evalues of 0 were equated with the maximum found evalue exponent plus 2, to give slightly more weight to identical proteins in contrast to minimally evolved proteins. Clustering was performed with the following algorithms: k-means (KMediansLloyd or KmeansMacQueen implementation), k-medians (KMediansLloyd implementation), k-medoids (CLARANS implementation), DBSCAN, EriC and meanshift. The value of k/epsilon/kernel-bandwidth was set between 2 to 50, besides for dbscan and meanshift, where the maximum was varied until all the sequences were merged into a single cluster. The value of minpts was set to 3 in all applicable cases. As initialization kmeansplusplus was used, where applicable, and at least two different runs were executed. All built in methods for cluster evaluation (e.g. Davies-Bouldin or silhouette-index (4-5)) were calculated for all clustering runs. The GH55, GH68, SFLD families 19 and 159 were investigated with clustering.

GNU parallel version 20161222 has been used during clustering and other steps of this research (6).

##### Network Clustering

Network clustering was performed with algorithms from the cdlib library, version 0.2.4 (7). The following 19 algorithms were used: belief, Chinese whispers, Eigenvector, GA, gdmp2, Girvan Newman (with parameter level=3), greedy modularity, Infomap, label propagation, Leiden, Louvain , lswl plus, Markov clustering, rber_pots, rb_pots, spinglass, surprise, threshold clustering and walktrap. All algorithms were used with standard parameters, unless mentioned. Evalues were converted to weights in the same way as for the clustering, using the exponent as weight. These algorithms were only tested on datasets GH51, GH54, GH55, GH68 and PL26. Artificial data was not used, since simulating a realistic weight distribution was infeasible.

##### Dimensionality reduction techniques

The same matrix which was used for the clustering was also used for the dimensionality reduction techniques. All methods (PCA, t-SNE, LLE, MDS, LTSA, Isomap and spectral embedding) were calculated in Python3 with Matplotlib v 3.1.3, Numpy v 1.17.2 and sklearn v 0.21.3 (8–11), based on an example on the scikit website by Jake Vanderplas. Fifty neighbors were used for the computations. T-SNE was run twice, once with PCA initialization, once with random initialization. Colouring was performed based on previously or newly assigned families, if available, to inspect these assignments. For PCA the loading of the components was investigated for an information gain as well. The networks for GH55, GH68, SFLD family 19 and 159 were investigated.

#### Supplementary Results

##### Automatic subfamily HMM building aids future annotations

To utilize these newly annotated subfamilies for future annotations, we decided to build HMM models for each subfamily, if at least 10 sequences were assigned at the selected subfamily level. Since this can reach a high number of subfamilies, as we have previously seen with GH16, and it would also already mean 12 subfamilies for GH51, we built these semi-automatically (see Materials and Methods).

In case of GH55, the chosen 3-subfamily scheme (SSN with $10^{-29}$ cutoff) leads to HMMs that assigned the correct subfamily in all cases, independent of any e-value threshold considerations.

In case of GH68, the chosen 2-subfamily scheme (SSN with $10^{-51}$ cutoff) lead to HMMs that resulted in at maximum 22 false positive (FP) and false negatives (FN) out of 1510 cases, if a minimum e-value of 10E-55 was used for assignment, with a minimum distance of 10E-55 to the next best subfamily HMM. For lower thresholds, mostly 3-9 FP and FNs emerged. Overall the best F1 value (defined as TP/(TP+0.5*(FP+FN)) could be seen if an e-value threshold between 10E-5 and 10E-35 was used, and no minimum difference or a minimum e-value difference of 10E-5 to the next best hit was used (see supplementary figure S4).

In case of GH51 had a total of 12 possible subfamilies, if any group with at minimum 10 sequences was considered (SSN with $10^{-58}$ cutoff). Of its 10,085 sequences, 10,001 are assigned to a subfamily, as defined by the SSN, and 84 sequences were present in groups smaller than 10 or as singles. This makes the choice of threshold more important, since in this case a big group of “true negative” (TN) assignments is necessary. The analysis shows that these TNs are causing the most incorrect assignments if low e-values were used, and that the TP sequences of the subfamilies were nearly always correctly assigned. The best tradeoff was seen if a minimum e-value of $10^{-45}$ was used, together with a minimum distance of $10^{-10}$ to the second best identification, resulting in 36 FP and 9 FN (together 0.45%). Even the worst considered option (E-value of $10^{-55}$, with $10^{-55}$ distance to the second best hit) yielded only 3% false assignments though, with most options generation <1% false assignments (supplementary figure S5).

**Clustering does not reach the same precision as SSNs**

Clustering is one of the simplest methods to assign groups in complicated data, and has been used since the middle of the last century (10). The different developed methods have different strengths and weaknesses, and their suitability cannot necessarily predicted from the given data. Since creating large amounts of SSNs can be demanding, computation and storage wise, we evaluated if clustering could maybe aid this effort. While we hypothesized that a centroid-based algorithm like k-means would give us suitable results, the lack of a noise concept was expected to be an issue, given that we also expect in some cases many singlets, which cannot be assigned to a cluster. We also hypothesized that some of the algorithms (e.g. correlation clustering like EriC) would find mathematically valid results, which we might not be able to interpret. We therefore parsed our results of the blast search into matrices instead of SSNs, with similarities based on absolute evalue exponents (e.g. 3.2E-20 would be represented as 20), and executed various clustering methods and evaluation criteria on these matrices.

Indeed, the clustering produced various outcomes, and indeed the centroid-based methods performed exactly according to our expectations. Results for kmeans, kmedians and kmedoids were evaluated based on Davies-Bouldin index, Gamma, Tau, C index and Calinski-Harabasz index, since these showed a non-monotonous increasing or decreasing pattern in at least some of the clustering runs. 10 different runs of these algorithms were investigated, due to the inherent stochasticity of these algorithms, which requires that the average of multiple results is considered. In case of GH55, the Davies-Bouldin index (4) often, but not always, identified 3 clusters as best for Kmeans/Kmedians/kmedoids, and these nearly perfectly corresponded to the 3 identified clusters in the SSNs (plus one single). The single seemed to disturb the assignments, since it also caused another sequence from a different cluster being assigned to the same cluster as itself.

In case of GH68, the centroid-based methods identified 5 clusters as being optimal (again based on the Davies-Bouldin index). Based on the SSN, our best split would only consist out of 2 subfamilies, but the next best split would create 4 main groups, so we compared the clustering results to these. Indeed, the clustering would retrieve 3 clusters nearly perfectly, would split the 4$^{th}$ into 2, and produce a composite cluster with the remainder and the smaller groups/singles.

The dbscan results were evaluated with the Davies-Bouldin index (after discarding all results where the algorithm had produced a single cluster, and any results which included >33% noise). For GH55, this nearly reproduced the same results as the centroid-based methods, although an incorrect sequence was assigned to noise. For GH68, dbscan also recovered the 4 previously identified clusters perfectly, and produced a 5$^{th}$ composite cluster from the remaining sequences. As second best result here a split into 2 clusters was defined, although this did not match the 2 clusters identified by the SSN approach. The Meanshift algorithm (evaluated with Davies-Bouldin index and Tau) produced similar results, although the evaluation with Tau for GH68 pointed to the 2 subfamily split, the evaluation with Davies-Bouldin pointed to the 4 subfamily split.

One algorithm, EriC, had a too long run time on even a small network like the GH55 or GH68 that we did not consider it reasonably usable for our purpose.

An attempt to cluster the SFLD families 19 and 159 was also performed, but a run time off weeks for all possible clustering runs turned out to be prohibitive.

**Network clustering does not always find appropriate divisions**

We first evaluated the network clustering results for GH55, since the division into three networks and one single node was hypothesized to not be challenging. Of the 19 utilized algorithms, four divided this network into three communities, one into four. One algorithm failed, whereas from the rest, all besides one divided this network into two communities. Of the algorithms predicting three communities, two (Chinese whispers and Markov clustering) predicted the three main groups correctly, and added the single sequence to one of these groups. The Infomap algorithm correctly predicted all four groups.

For GH68, 18 out of the 19 algorithms produced a result, and one aborted due to insufficient memory. Six algorithms predicted no communities, nine predicted two, the remaining four algorithms predicted three or four communities. Of the nine algorithms predicting two communities, four (including Chinese whispers, Markov clustering and Infomap) predicted the two communities nearly perfect, with assigning between three to five sequences at the interface between both clusters to the wrong cluster. These wrongly assigned sequences corresponded to the same sequences which produced false-positives in the HMM investigation. Of the algorithms predicting three to four sequences, none predicted the grouping at the levels of $10^{-51}$ or $10^{-113}$, although the threshold clustering algorithm predicted an intermediate state.

For GH51, one algorithm failed due to memory requirements, two algorithms did not predict any communities, and another two separated each sequence into its own community (predicting 10086 communities). The remaining algorithms predicted between two to 28 communities. Of these, five separated the two major groups into two different groups, but failed at the finer resolution, and one separated the three major groups, but didn’t separate the smaller groups either. The remaining eight algorithms did not result in any prediction coherent with any investigated network threshold.

For GH54, for which we did not predict any subfamilies based on the SSN, 18 out of 19 algorithms finished their prediction. Of these twelve predicted only a single network. Of the remaining six, two predicted two networks of which one consisted only out of a single sequence. The remaining four algorithms did not predict any grouping which would agree with any SSN prediction.

For PL26, for which we predicted an optimal grouping of one main network and one smaller network of 21 nodes, all 19 algorithms finished processing. Twelve predicted a single network. Of the remaining seven, only one (threshold clustering) correctly predicted the group of 21 nodes, but also predicted a second group. The six other algorithms did not yield any results which were in agreement with any of the SSN predictions.

#### Dimensionality reductions techniques show grouping in a non-distinguishable  way

Dimensionality reduction techniques like PCA or the recently developed t-SNE are methods used in the scientific community to estimate groupings in their data. E.g. t-SNE is one of the most widely used techniques in single-cell data to visualize different cell populations (12), and PCA is known to most researchers, independent of field. We therefore assumed that these methods should be able to derive the subgrouping without actually constructing any SSNs. We performed PCA, t-SNE and various other techniques on the same matrices used as input for the clustering, and visually investigated the 2-dimensional output, as well as the loading of the components. 3 of the techniques often either failed on the big amount of data or compressed the results into too little points (LLE, LTSA, SE). Isomap, PCA and t-SNE with PCA initialization had in some cases as result a star-like formation with 3 or 4 arms, with most of the data being continuously distributed, without a clear grouping. While the coloring of the previously determined subgroups showed that they were indeed grouped together, it was not possible to discriminate between subgroups or to estimate the possible amount of groups, unless very little, simple data was used, as in the case for GH55 (see supplementary figure S6, PCA of  GH55, GH68, SFLD 159 and SFLD 19). MDS showed a distinctive pattern, also grouping families mostly together, but did not allow for discrimination either. t-SNE with random initialization seemed to show a more random result, although sometimes families were grouped together.

From these results, we concluded that these technologies were suitable for our purpose.

#### Supplementary Figures

Figure S1: A) One network set, consisting out of two networks, which are connected with increasing amounts of edges. B) Special case #1: one network is connected to a network of singles. Each single node is consisted first by only one connection, and not random, preventing a second connection before all nodes are connected. C) Special case #2: one network is connected to a network of singles. Each single node is connected first to all nodes in the main network.

Figure S2: Behaviour of different metrics over two networks the size of 50. A) Closeness and degree centrality both favour more edges over less, and both networks combined with zero connections score worse than a network with one connection. The adjustment to score these networks better has been performed in this work. B) The number of edges, the s-metric, the average degree, subgraph centrality and harmonic centrality show a monotonous behaviour. The more edges a component possesses, the better the score. This means that one perfect network scores worse than two perfect networks with one connection. C) Betweenness and similar metrics score in an inverted way to closeness in A). Based on this behaviour it cannot be decided which metric is more suitable. A decision had to be made based on the performance with single nodes, as discussed in the manuscript. D) The amount of bridges, components and communities does not show a quantitative enough behaviour. Many instances are scored with a value of one, making it not possible to discriminate between better or worse networks. E) and F) show the behaviour of the remaining four criteria. The eigenvector centrality had to be plotted separately, without the first three data points, due to differences in scale between these data points. All four metrics show a behaviour which might favour less edges over more edges at some point in the data set, making their behaviour not desirable. Parameters may have been scaled for display purposes.

Figure S3: Violin plot of percentage identity values matched with corresponding evalue values for PL26. With 32% identity between two sequences the first evalue of zero was obtained. With 64% identity between two sequences all evalues correspond to zero.

Figure S4: F1 score of the HMM subfamily assignments for GH68. 2 new subfamily HMMs were built, and the best F1 score with least FP and FN was seen if a new sequence was assigned to a subfamily if the HMM reported an e-value between $10^{-5}$ and $10^{-35}$. The distance to the next best HMM hit was also considered, but choosing no difference (first line in legend) or choosing a minimum of $10^{-5}$ e-value difference to the second best hit (second line in legend) did not make any difference in this case (graphs overlap).

Figure S5: F1 score of the HMM subfamily assignments for GH51. 12 new subfamily HMMs were built, and the best F1 score with least FP and FN was seen if a new sequence was assigned to a subfamily if the HMM reported an e-value at 10E-45. In addition, the distance to the next best HMM hit has to be considered, since requiring a minimum of 10E-10 between the best HMM hit and the second best HMM hit gave the best result.

Figure S6: Example of dimensionality reduction results. Colouring according to groups identified by the SSNs analysis. For GH68, both groupings at 10E-51 and 10E-154 are displayed. For the SFLD families, pre-computed groups were used. While in the PCA plot for GH55 the three subfamilies are visible, this gets ambigious in the MDS plot or for GH68, and indistinguishable for the SFLD families.

### References


1. Viborg AH, Terrapon N, Lombard V, Michel G, Czjzek M, Henrissat B, et al. A subfamily roadmap of the evolutionarily diverse glycoside hydrolase family 16 (GH16). J Biol Chem. 2019 Nov;294(44):15973–86.

2. Sievers F, Higgins DG. Clustal Omega for making accurate alignments of many protein sequences: Clustal Omega for Many Protein Sequences. Protein Sci. 2018 Jan;27(1):135–45.

3. Macqueen J. Some methods for classification and analysis of multivariate observations. In: Proceedings of the 5th Berkeley Symposium on Mathematical Statistics and Probability. p. 281–97.

4. D. L. Davies, D. W. Bouldin. A Cluster Separation Measure. IEEE Transactions on Pattern Analysis and Machine Intelligence. 1979 Apr;PAMI-1(2):224–7.

5. Schubert E, Zimek A. ELKI: A large open-source library for data analysis - ELKI Release 0.7.5 “Heidelberg.” arXiv:190203616 [Internet]. 2019 [cited 2021 Jun 25]; Available from: http://arxiv.org/abs/1902.03616

6. Tange O. GNU Parallel: The Command-Line Power Tool. ;login. 2011;36(1):42–7.

7. Rossetti G, Milli L, Cazabet R. CDlib: a Python Library to Extract, Compare and Evaluate Communities from Complex Networks. Applied Network Science Journal. 2019 4(25)

8. Harris CR, Millman KJ, van der Walt SJ, Gommers R, Virtanen P, Cournapeau D, et al. Array programming with NumPy. Nature. 2020 Sep 17;585(7825):357–62.

9. Pedregosa F, Varoquaux G, Gramfort A, Michel V, Thirion B, Grisel O, et al. Scikit-learn: Machine Learning in Python. J Mach Learn Res. 2011;12:2825–30.

10. SciPy 1.0 Contributors, Virtanen P, Gommers R, Oliphant TE, Haberland M, Reddy T, et al. SciPy 1.0: fundamental algorithms for scientific computing in Python. Nat Methods. 2020 Mar;17(3):261–72.

11. J. D. Hunter. Matplotlib: A 2D Graphics Environment. Computing in Science & Engineering. 2007 Jun;9(3):90–5.

12. Kobak D, Berens P. The art of using t-SNE for single-cell transcriptomics. Nat Commun. 2019 Dec;10(1):5416.
